# A comparative atlas of single-cell chromatin accessibility in the human brain

**DOI:** 10.1101/2022.11.09.515833

**Authors:** Yang Eric Li, Sebastian Preissl, Michael Miller, Nicholas D. Johnson, Zihan Wang, Henry Jiao, Chenxu Zhu, Zhaoning Wang, Yang Xie, Olivier Poirion, Colin Kern, Antonio Pinto-Duarte, Wei Tian, Kimberly Siletti, Nora Emerson, Julia Osteen, Jacinta Lucero, Lin Lin, Qian Yang, Quan Zhu, Sarah Espinoza, Anna Marie Yanny, Julie Nyhus, Nick Dee, Tamara Casper, Nadiya Shapovalova, Daniel Hirschstein, Rebecca D. Hodge, Sten Linnarsson, Trygve Bakken, Boaz Levi, C. Dirk Keene, Jingbo Shang, Ed S. Lein, Allen Wang, M. Margarita Behrens, Joseph R. Ecker, Bing Ren

**Affiliations:** Department of Cellular and Molecular Medicine, University of California, San Diego, San Diego, CA, USA; Center for Epigenomics, University of California San Diego, School of Medicine, La Jolla, CA, USA.; The Salk Institute for Biological Studies, La Jolla, CA, USA; Department of Computer Science and Engineering, University of California San Diego, CA, USA; Division of Molecular Neurobiology, Department of Medical Biochemistry and Biophysics, Karolinska Institute, Stockholm, Sweden; Allen Institute for Brain Science, Seattle, WA, USA; Department of Laboratory Medicine and Pathology, University of Washington, Seattle, WA, USA; Howard Hughes Medical Institute, The Salk Institute for Biological Studies, La Jolla, CA, USA; Ludwig Institute for Cancer Research, La Jolla, CA, USA

## Abstract

The human brain contains an extraordinarily diverse set of neuronal and glial cell types. Recent advances in single cell transcriptomics have begun to delineate the cellular heterogeneity in different brain regions, but the transcriptional regulatory programs responsible for the identity and function of each brain cell type remain to be defined. Here, we carried out single nucleus ATAC-seq analysis to probe the open chromatin landscape from over 1.1 million cells in 42 brain regions of three neurotypical adult donors. Integrative analysis of the resulting data identified 107 distinct cell types and revealed the cell-type-specific usage of 544,735 candidate cis-regulatory DNA elements (cCREs) in the human genome. Nearly 1/3 of them displayed sequence conservation as well as chromatin accessibility in the mouse brain. On the other hand, nearly 40% cCREs were human specific, with chromatin accessibility associated with species-restricted gene expression. Interestingly, these human specific cCREs were enriched for distinct families of retrotransposable elements, which displayed cell-type-specific chromatin accessibility. We uncovered strong associations between specific brain cell types and neuropsychiatric disorders. We futher developed deep learning models to predict regulatory function of non-coding disease risk variants.

## Main Text

Neurological disorders and mental illnesses are the leading cause of disease burdens in the United States(*1*). Tens of thousands of sequence variants in the human genome have been linked to the etiology of neuropsychiatric disorders(*2, 3*). However, interpreting the mode of action of the identified risk variants remains a daunting challenge since the vast majority of them are non-protein-coding (*4, 5*). It is increasingly suspected that a large fraction of the non-coding risk variants contribute to disease etiology by perturbing transcriptional regulatory elements and target gene expression in the disease-relevant cell types(*6–8*). However, a lack of the maps and tools to explore gene activities and their transcriptional regulatory sequences at high cellular and anatomical resolution in the brain prevents a clearer mechanistic understanding of the broad spectrum of neuropsychiatric disorders.

The human brain is made up of hundreds of billions of neurons, which through trillions of synapses form a complex neurocircuitry to carry out diverse neurocognitive functions. The functionality of the neural circuitry is supported and maintained by an even greater number of glial cells including astrocytes, oligodendrocytes, oligodendrocyte precursor cells, and microglia, among others. Single-cell RNA-seq and high throughput imaging experiments have produced detailed cell taxonomies for mouse brains and a few cortical regions in human (*9–17*), leading to a comprehensive view of cell types and their molecular signatures in several brain regions(*18–20*). Analysis of gene expression patterns using single-cell transcriptomics and spatial transcriptomics assays (*9, 11, 12, 16, 21–24*), have further advanced our understanding of the transcriptional landscapes in different brain cell types. In contrast to the large body of transcriptomic analyses, analysis of the regulatory elements that drive the cell-type specific expression of genes is lagging. Current catalogs of candidate regulatory sequences in the human genome, most notably those generated by ENCODE and Epigenome Roadmap consortia (*6–8, 25, 26*), still lack the information about cell-type-specific activities of each element especially those identified from brain tissues, because conventional assays performed using bulk tissue samples, unfortunately, fail to resolve cCREs in individual cell types comprising the heterogeneous tissues. To address this problem, recent technological advances have enabled the analysis of open chromatin at single cell resolution (*27–32*) in adult mouse tissues and several brain regions (*27, 29, 33–35*), generating cell-type-specific maps of gene regulatory elements for a limited number of human brain cell types and brain regions (*36–38*).

As part of the BRAIN Initiative Cell Census Network (BICCN), we have carried out single-cell profiling of transcriptome, chromatin accessibility, and DNA methylome across >40 regions in the human brain from multiple neurotypical adult donors. Here, we describe a single-cell chromatin accessibility atlas comprising ∼1.1 million human brain cells and results from integrative analysis of this genomic resource. We used this chromatin atlas to define 107 distinct cell types, and uncover the state of chromatin accessibility at 544,735 cCREs in one or more of these brain cell types. We integrate our chromatin atlas with the companion single cell transcriptome and DNA methylome atlases to link cCREs to putative target genes. We compared with a previous catalog of cCREs from mouse cerebrum, finding strong evolutionary conservation in both sequence and chromatin accessibility for nearly 1/3 of human brain cCREs. We further predict disease relevant cell types for 19 neurological traits and diseases. Finally, we developed machine learning models to predict the regulatory function of disease risk variants. We created an interactive web atlas to disseminate this resource (cis-element ATLAS [CATLAS]; http://catlas.org).

## Result

### A single-cell chromatin accessibility atlas of human brains

We dissected 42 brain regions from the human cortex (CTX), hippocampus (HIP), basal nuclei (BN), midbrain (MB), thalamus (THM), cerebellum, and pons from three neurotypical male donors (D1, D2, and D4) at the age of 29, 42, and 58, respectively (**Fig. 1A, Table S1**), according to the Allen Brain Reference Atlas (*39*). For each brain sample, we performed snATAC-seq using a protocol described previously (*40*) (**Fig. 1A, Fig. S1A-D, Table S2**). The reliability of the data was evidenced by sequencing reads showing nucleosome-like periodicity (**Fig. S1E**), excellent correlation between datasets from similar brain regions across the three donors (**Fig. S1F**), high transcription start site (TSS) enrichments as well as other quality control metrics (see **Methods**). A total of 1,290,974 nuclei passed stringent quality control criteria (**Fig. S1G**, see **Methods**). After removing an additional 156,614 snATAC-seq profiles that likely resulted from potential barcode collision or doublets (**Fig. S1H-J**, see **Methods**), a total of 1,134,360 nuclei were retained. Among them, 595,713 were from cortex, 72,190 from HIP, 317,480 from BN, 23,114 from MB, 50,768 from THM, 51,775 cerebellum and 25,459 from pons (**Table S3**). On average 4,970 chromatin fragments were detected in each nucleus (**Table S3, Fig. S1K-M**, see **Methods**).

**Fig. 1.**
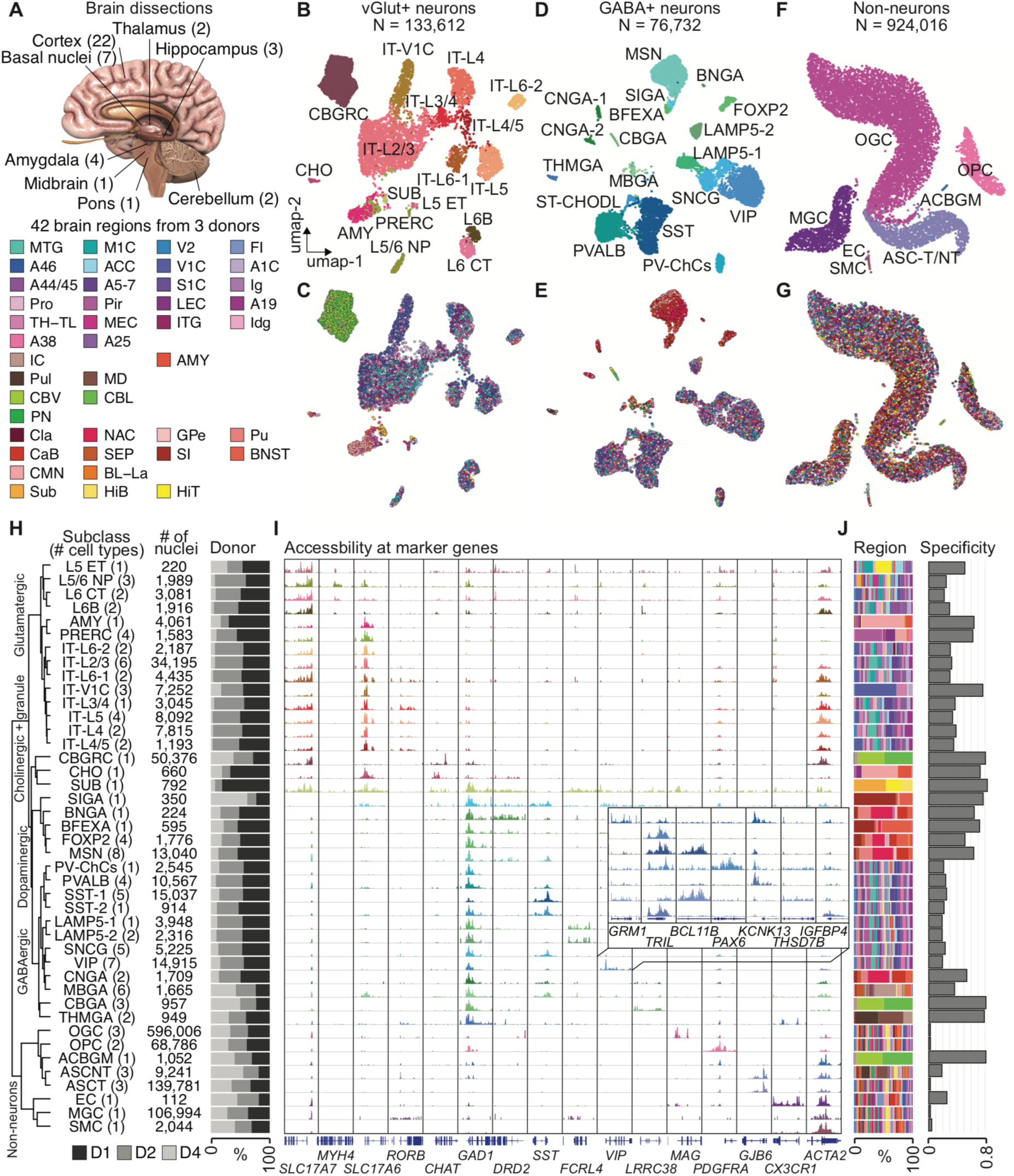
Single-cell analysis of chromatin accessibility in the human brain. (**A**) Schematic of sampling strategy and brain dissections. A detailed list of regions is provided in **Table S1**. (**B**) Uniform manifold approximation and projection (UMAP) embedding and clustering analysis of glutamatergic neurons from snATAC-seq data. Individual nuclei are colored by cell subclasses. A full list and description of cluster labels are provided in **Tables S3**. (**C**) UMAP embedding of glutamatergic neurons, colored by brain regions. (**D**) UMAP embedding and clustering analysis of GABAergic neurons, colored by cell subclasses. (**E**) UMAP embedding of GABAergic neurons, colored by brain regions. (**F**) UMAP embedding and clustering analysis of non-neurons, colored by cell subclasses. (**G**) UMAP embedding of non-neurons, colored by brain regions. (**H**) Left, hierarchical organization of 42 cell subclasses on chromatin accessibility. Middle, the number of nuclei in each subclass. Right, Bar chart representing the relative contribution of 3 donors to each cell subclass. (**I**) Genome browser tracks of aggregate chromatin accessibility profiles for each subclass at selected marker gene loci that were used for cell cluster annotation. A full list and description of subclass annotations are in **Table S4**. (**J**) Left, bar chart representing the relative contribution of brain regions to cell subclasses. Right, regional specificity scores of cell subclasses.

We next carried out iterative clustering with the above snATAC-seq profiles and classified them into three major classes: glutamatergic (vGlut+) or excitatory neurons (11.8%), GABAergic (GABA+) or inhibitory neurons (6.8%) and non-neuronal cells (81.4%) (**Fig. 1B, D, F, Fig. S2, Fig. S3A-B**). We further classified the three cell classes into 14 sub-classes of glutamatergic neurons, 2 sub-classes of granule cell types, 1 sub-class of cholinergic neurons, 4 sub-classes of dopaminergic neurons, 2 sub-classes of thalamic and midbrain derived neurons, 11 sub-classes of cortical GABAergic neurons, and 8 sub-classes of non-neuronal cells (**Fig 1B, D, F**). Each sub-class was annotated based on chromatin accessibility at promoters and gene bodies of at least three marker genes of known brain cell types, together with the brain region (**Fig. 1C, E, G, Fig. S3C, Table S4, and S5**). For each sub-class, we also conducted a third round of clustering and identified a total of 107 distinct brain cell types (**Fig. 1H, Fig. S4, Table S3**, see **Methods**). To determine the optimal number of cell types within each sub-class, we evaluated the relative stability from a consensus matrix based on 100 rounds of clustering with randomized starting seeds (see **Methods**). We then calculated the proportion of ambiguous clustering (PAC) score and dispersion coefficient (DC) to find the optimal resolution (local minimum and maximum) for cell type clustering (**Fig. S2B-E**). For example, *VIP* positive (*VIP^+^*) neurons (VIP) were further divided into multiple cell types with distinct chromatin accessibility at multiple gene loci (**Fig. 1I**, **Fig. S2B-E, Fig. S4**). We found that the clustering result of snATAC-seq was robust to variation of sequencing depth, and signal-to-noise ratios, and most cell sub-classes showed no batch effect from at least two donors using local inverse Simpson’s index (LISI) analysis, with the exception of two cell sub-classes (SUB, granule cells from hippocampus, SMC, vascular smooth muscle cells) that were mostly captured from one donor (**Fig. S5**). To capture the relative similarity in chromatin landscapes among the 42 sub-classes we constructed a robust hierarchical dendrogram (**Fig. 1H**, **Fig. S7**). This dendrogram shows known organizing principles of human brain cells: the non-neuronal class is separated from the neuronal class, which are further separated based on neurotransmitter types (GABAergic, dopaminergic, cholinergic, and glutamatergic) and developmental origins (**Fig. 1H**, **Fig. S6**, see **Methods**).

All neuronal and some glial cell types are distributed in the human brain in a non-uniform fashion (**Fig. 1J**). We defined a regional specificity score for each sub-class based on the contribution from different brain regions. While the majority of glial cell types are ubiquitously distributed throughout the brain, showing very low regional specificity (**Fig. 1J**, right), there are exceptions. For example, the Bergmann glia (ACBGM) is specifically found in the cerebellum. On the other hand, all neuron types are characterized by moderate to high regional specificity (**Fig. 1J**, right). We found a stark separation based on brain sub-regions for distinct neuron types including the granular cells in the cerebellum (CBGRC) and medium spiny neurons (MSN) in the basal ganglia. For glutamatergic neurons, we also observed distinct types of intra-telencephalic (IT) cortical neurons in the primary visual cortex (V1C) (IT-V1C), and excitatory neurons from the amygdala (AMY) to be highly restricted to specific brain regions or dissections.

We also compared our single-nucleus chromatin accessibility derived cell clusters with the cell taxonomy defined from other modalities. We performed the integrative analysis with single-cell RNA-seq (scRNA-seq) and single-nucleus methylome sequencing (snmC-seq) (companion papers, Siletti, et al. 2022 and Tian, et al. 2022, see **Methods**). For 38 of 42 sub-classes defined based on chromatin accessibility (A-Type), we could identify a corresponding sub-class defined based on transcriptomics (T-Type; overlap score cut-off >0.2, **Fig. S7**). In addition, for 39 out of 42 A-types we identified one, or in some cases more, corresponding sub-class defined based on DNA methylation (M-Type, **Fig. S7**). Of note, a few cell clusters can hardly be aligned with other modalities, possibly due to differences in brain dissections and/or under sampling of neurons in our datasets. For example, the A-type granule cells from cerebellum (CBGRC) were not identified in T-types, but aligned well with M-type. The cholinergic neurons were not captured in M-types, but aligned to a mixture class of T-types called Glycinergic, cholinergic, monoaminergic, or peptidergic neurons.

### Mapping and characterization of human brain cCREs

As a first step towards defining the gene regulatory programs that underlie the identity and function of each brain cell type, we identified the open chromatin and candidate *cis-regulatory* DNA elements (cCREs) in each of the 107 brain cell types. We aggregated the chromatin accessibility profiles from the nuclei comprising each cell cluster/type and identified the open chromatin regions with MACS2(*41*) (**Fig. S8A**). We filtered the resulting accessible chromatin regions based on whether they were called in at least two donors, or in two pseudo-replicates (**Fig. S8A-B**). From our previous study, we found that read depth or cluster size can affect MACS2 peak calling scores. We therefore used “score per million” (SPM)(*42*) to correct this bias (**Fig. S8C**, see **Methods**). About 1000 nuclei in a cluster were sufficient to identify over 80% of the accessible regions in a cell type, consistent with our previous finding(*40*) (**Fig. S8C**). We iteratively merged the open chromatin regions identified from every cell type, and kept the summits with the highest MACS2 score for overlapped regions. On average, we detected 62,045 open chromatin regions per cell type (500 bp in length), and a union of 544,735 open chromatin regions across all 107 cell types (**Fig. S8D, Table S6, and S7,** see **Methods**). These cCREs together made up 8.8% of the human genome (hg38) (**Table S8**). Of these cCREs, 95.3% were located at least 2 kbp away from annotated promoter regions of protein-coding and lncRNA genes (**Fig. 2A**, **Table S8**). The promoter-distal cCREs were distributed in introns (34.8%), intergenic (27.8%), and other genomic regions. Interestingly, 22% of them overlap with endogenous retrotransposable elements, including long terminal repeats class (LTR, 6.8%), LINEs (long interspersed nuclear elements, 11.3%), SINEs (short interspersed nuclear elements, 3.9%). Several lines of evidence support the authenticity of the identified cCREs. First, both proximal and distal cCREs showed higher levels of sequence conservation than random genomic regions with similar GC content (**Fig. 2B**). Second, 89.6% of cCREs overlapped with DNase hypersensitive sites (DHS) previously mapped in a broad spectrum of bulk human tissues and cell types including fetal and adult brains (*43*). This list further expands candidate CREs previously annotated in the human genome by the ENCODE(*26*) and a recent survey of chromatin accessibility in single nuclei across fetal and adult human tissues (*44*)(**Fig. 2C**).

**Fig. 2:**
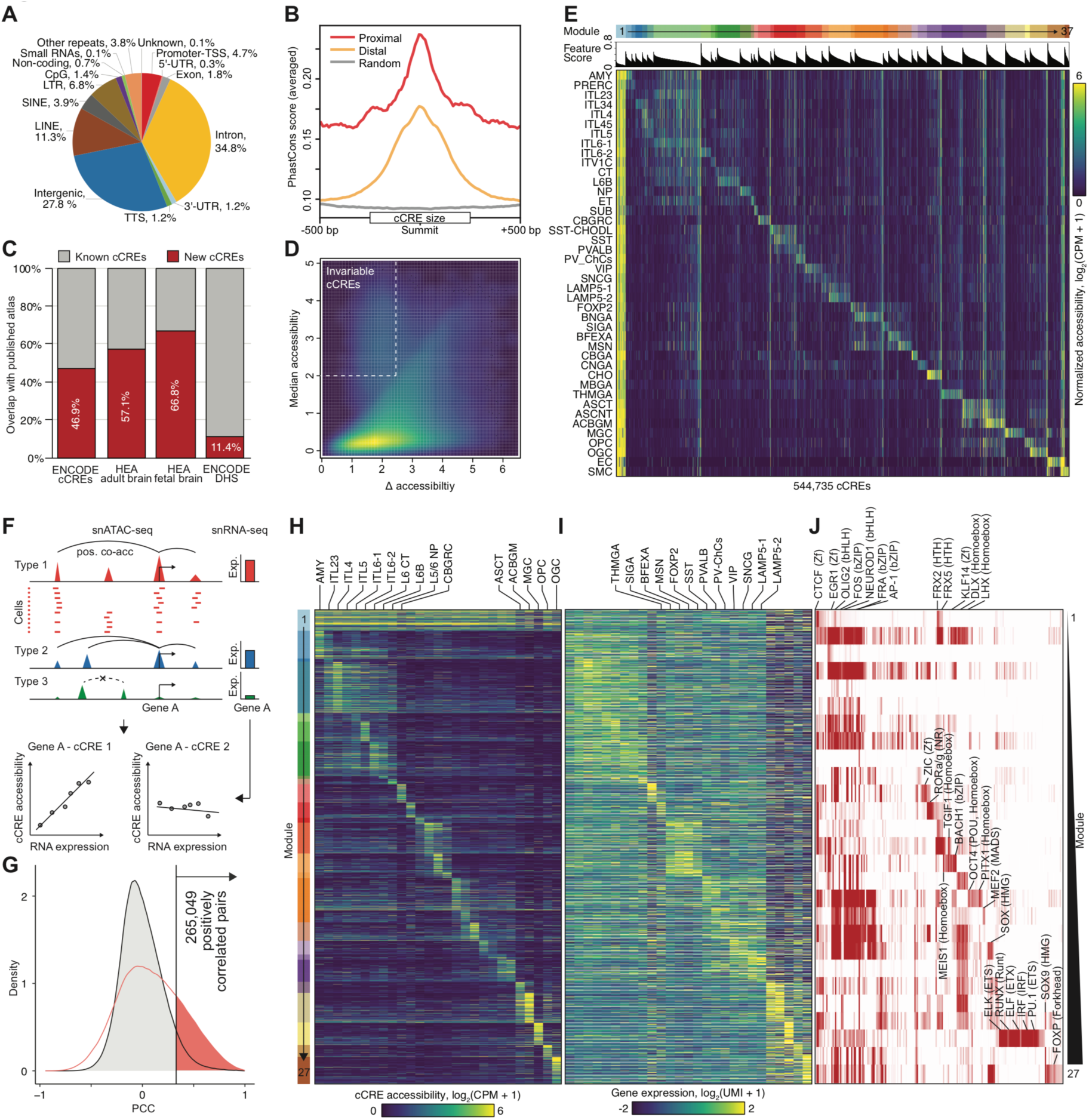
Identification and characterization of candidate CREs (cCREs) across human brain cell types. (**A**) Pie chart showing the fraction of cCREs that overlaps with different classes of annotated sequences in the human genome. TSS, transcription start site; TTS, transcription termination site. UTR, untranslated region. LINE, long interspersed nuclear element. SINE, short interspersed nuclear element. LTR, long terminal repeats. (**B**) Average phastCons conservation scores of proximal (in red) and distal cCREs (in yellow), and random genomic background is indicated in gray. (**C**) Stacked bar plot showing the percentage of new cCREs defined in this study (in red) and percentage of cCREs that overlapped with public recourse (in grey), including the cCREs and DHSs in the SCREEN database, cCREs identify in human enhancer atlas (HEA) fetal and adult brain. (**D**) Density map comparing the median and maximum variation of chromatin accessibility at each cCRE across cell types. Each dot represents a cCRE. (**E**) Heat map showing association of the 42 subclasses (rows) with 37 cis-regulatory modules (top, from left to right). Columns represent cCREs. A full list of subclass or module associations is in **Table S9**, and the association of cCREs to modules is in **Table S10**. CPM, counts per million. (**F**) Schematic overview of the computational strategy used to identify cCREs that are positively correlated with transcription of target genes. (**G**) In total, 265,049 pairs of positively correlated cCRE and genes (highlighted in red) were identified (FDR < 0.01). Grey filled curve shows distribution of Pearson’s correlation coefficient (PCC) for randomly shuffled cCRE–gene pairs. (**H**) Heat map showing chromatin accessibility of putative enhancers and (**I**) the expression of linked genes (right). Genes are shown for each putative enhancer separately. UMI, unique molecular identifier. (**J**) Enrichment of HOMER known transcription factor (TF) motifs in distinct enhancer–gene modules.

To define the cell type specificity of the above cCREs, we first plotted the median levels of chromatin accessibility against the maximum variation for each element (**Fig. 2D**). We found that the majority of cCREs displayed highly variable chromatin accessibility across the brain cell types identified in the current study, except for a small proportion of invariable cCREs (2.0 %) that showed accessibility in virtually all cell clusters, of which 87% were at proximal regions to TSS (**Fig. 2D**). To characterize the cell type specificity of the cCREs more explicitly, we used non-negative matrix factorization to group them into 37 modules, with elements in each module sharing similar cell type specificity profiles. Except for the first module (M1) which included mostly cell-type invariant cCREs, the remaining 36 modules displayed highly cell-type restricted accessibility (**Fig. 2E**, **Table S9, and S10**). These cell type restricted cCRE modules were enriched for distinct sets of motifs recognized by known transcriptional regulators (**Table S11**). For example, the NEUROG2 and ASCL1 enriched in module M4 for intratelencephalic (IT) neurons at cortical layer 2/3 (IT-L2/3) have been reported to be proneural genes and critical for cortical development (**Table S11**)(*45*). The SOX family factors in module M35 for oligodendrocytes (OGC) are pivotal regulators of a variety of developmental processes (**Table S11**)(*46*). These results lay a foundation for dissecting the gene regulatory programs in different brain cell types and regions.

### Linking distal cCREs to target genes

To investigate the transcriptional regulatory programs that are responsible for cell-type-specific gene expression patterns in the human brain, we carried out an integrative analysis that combines the snATAC-seq data collected in the current study with scRNA-seq data generated by a companion paper (Siletti, et al. 2022) from matched brain regions (**Fig. S7**). We first connected 255,828 distal cCREs to 14,861 putative target genes by measuring the co-accessibility across single nuclei in every cell sub-class using Cicero(*47*) (**Fig. 2F**, upper, see **Methods**), which resulted in a total of 1,661,975 gene–cCRE pairs within 500 kb of each other. Next, we identified the subset of cCREs the accessibility of which positively correlates with the expression of putative target genes and therefore function as putative enhancers in neuronal or non-neuronal types (**Fig. 2F**, bottom). This analysis was restricted to distal cCREs and expressed genes captured from 27 matched cell sub-classes defined above from integrative analysis between snATAC-seq and scRNA-seq (**Fig. S7**). We revealed a total of 265,049 pairs of positively correlated cCRE (putative enhancers) and genes at an empirically defined significance threshold of FDR < 0.05 (**Table S12**). These included 114,877 putative enhancers and 13,094 genes (**Fig. 2G, Fig. S9, Table S12**). The median distance between the putative enhancers and the target promoters was 176,345 bp (**Fig. S9A**). Each promoter region was assigned to an average of 7 putative enhancers, and each putative enhancer was assigned to two genes on average (**Fig. S9B-C**).

We further classified these putative enhancers into 27 modules by using non-negative matrix factorization (**Table S13, and S14**), to investigate how cell-type-specific gene expression is regulated. The putative enhancers in each module displayed a similar pattern of chromatin accessibility across cell sub-classes (**Fig. 2H**), and the expression of putative target genes showed a correlated pattern (**Fig. 2I**). This analysis also revealed a large group of 5,113 putative enhancers that were linked to 4,775 genes expressed at a higher level across all neuronal cell clusters than in non-neuronal cell types (module M1) (**Fig. 2H, I, Table S13, and S14**). These putative enhancers are strongly enriched for CTCF, and RFX binding sites (**Table S15**), which is consistent with what we previously found in the mouse cerebrum(*40*).

We also uncovered modules of enhancer–gene pairs that were active in a more restricted manner (modules M2–M27) (**Fig. 2H-J**). For example, we identified modules (M2-M7) associated with several cortical glutamatergic neurons (IT-L2/3, IT-L4, IT-L5, IT-L6-1, IT-L6-2), in which the putative enhancers were enriched for sequence motifs recognized by the bHLH factors NEUROD1 (**Fig. 2J**, **Table S13-S15**). Another example was module M15 associated with medium spiny neurons (MSN), in which putative enhancers were enriched for motifs recognized by MEIS factors, which play an important role in establishing the striatal inhibitory neurons (**Fig. 2J**, **Table S13-S15**). Module M25 was associated with microglia (MGC). Genes linked to putative enhancers in this module were related to immune genes and the putative enhancers were enriched for the binding motif for ETS-factor PU.1, a known master transcriptional regulator of this cell lineage (**Fig. 2J**, **Table S13-S15**). This observation is consistent with the paradigm that cell-type-specific gene expression patterns are largely established by distal enhancer elements.

### Regional specificity of glial and neuronal cell cCREs

The single-cell atlas of chromatin accessibility generated in this study provides a unique opportunity to characterize the heterogeneity of the gene regulatory programs that might underlie the specialized functions of glial and neuronal cells in each brain region. While most non-neuronal cell types, including oligodendrocytes (OGCs), oligodendrocyte precursor cells (OPCs), microglia (MGC), telencephalon astrocytes (ASCTs), non-telencephalon (ASCNTs), and various vascular cells were ubiquitously distributed throughout the different brain dissections (**Fig. 1J**), molecular diversity has been recently reported in these cells in juvenile and adult vertebrates(*48–51*). We therefore leverage the largest collection of >900,000 single nuclei of non-neuronal cells and the high-resolution brain dissections in the present study (**Fig. S1N**), to more thoroughly characterize the regulatory diversity of the non-neuronal populations.

The UMAP embeddings in the brain regional spaces showed a gradient among cell types of OGCs, OPCs, MGCs, and ASCTs (**Fig. 3A-B, E-F, I-J, and M-N, Fig. S10**). We hypothesized that these gradients may reflect heterogeneity in cCRE usage in these glial cells across brain regions. We first calculated the averaged chromatin accessibility and coefficient of variation (CV) across 42 brain regions for every cCRE identified in OGCs, OPCs, MGCs, ASCTs, respectively (**Fig. 3C, G, K, and O, Fig. S10**). To our surprise, a large number of cCREs displayed highly variable chromatin accessibility across the brain regions (**Fig. 3C, G, K, and O, Fig. S10**). In total 55,304 or 40.1% of total cCREs identified in OGCs, 43,574 or 33.0% of total cCREs identified in OPCs, 37,962 or 34.5% of total cCREs identified in MGC, and 46,979 or 33.1% of total cCREs identified in ASCTs, were found to show variability in chromatin accessibility across major brain regions. These variable cCREs show similar regional specificities across three donors. Next, applying k-means clustering analyses to the variable cCREs identified in the above glial cell populations (**Fig. 3D, H, L, and P**), we revealed distinct open chromatin patterns in OGCs, OPCs, and ASCTs from the cerebellum (CB). Interestingly, a large fraction of these variable cCREs showed higher chromatin accessibility in the cerebellum (**Fig. 3D, H, and P**). We also observed loss of chromatin accessibility in a large number of cCREs in distinct brain structures (**Fig. 3D, H, L, and P**).

**Fig. 3:**
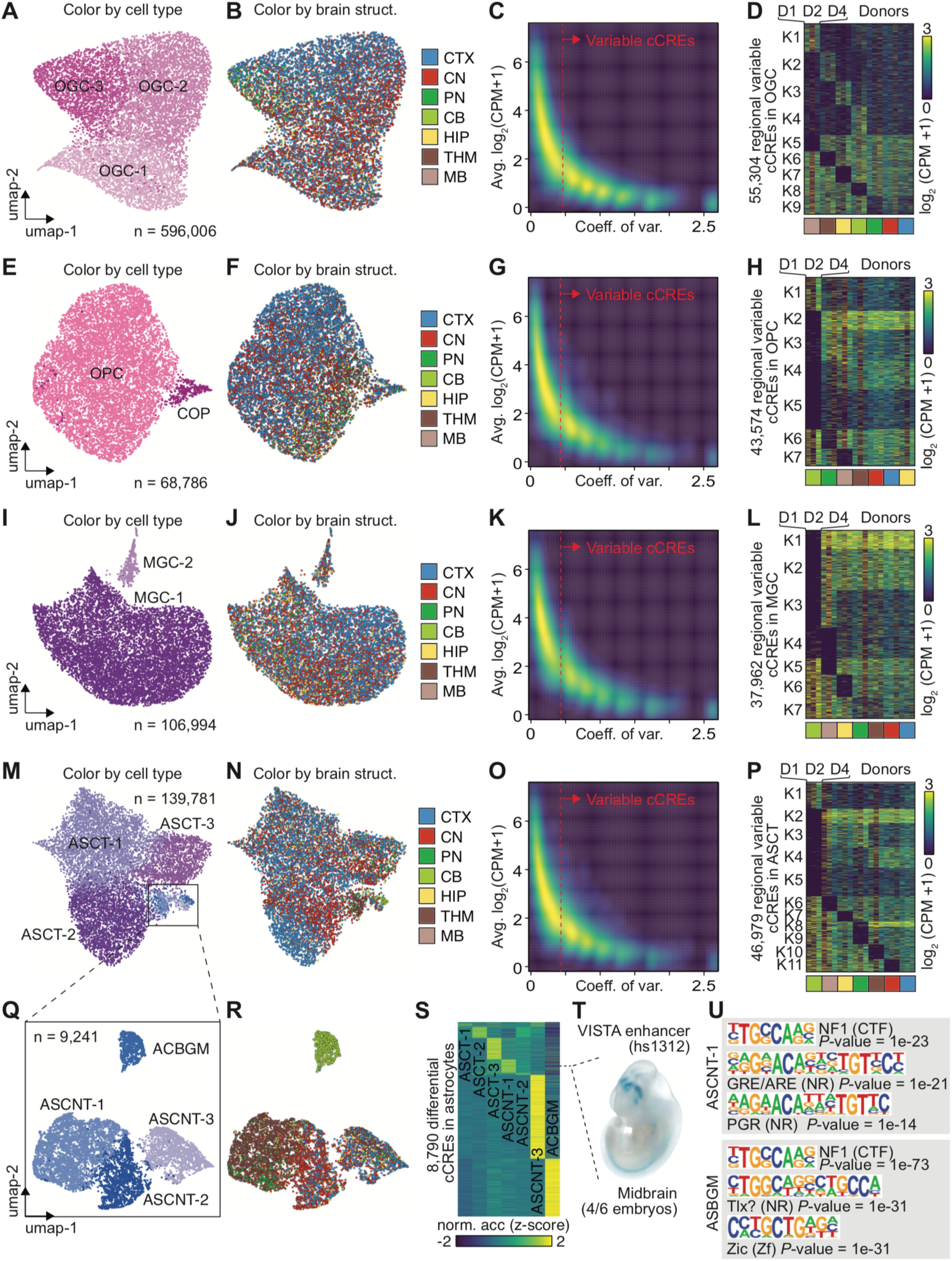
Regional specificity of cell types correlates with chromatin accessibility. (**A**) UMAP embedding of cell types of oligodendrocytes (OGCs). (**B**) UMAP embedding of oligodendrocytes, colored by major brain structures. (**C**) Density scatter plot comparing the averaged accessibility and coefficient of variation across brain structures at each cCRE. Variable cCREs for OGCs are defined on the right side of dash line. (**D**) Heat map showing the normalized accessibility of variable cCREs in OGCs. CPM, counts per million. (**E**) UMAP embedding of cell types of oligodendrocyte precursor cells (OPCs). (**F**) UMAP embedding of OPCs, colored by major brain structures. (**G**) Density scatter plot comparing the averaged accessibility and coefficient of variation across brain structures at each cCRE. Variable cCREs for OPCs are defined on the right side of dash line. (**H**) Heat map showing the normalized accessibility of variable cCREs in OPCs. (**I**) UMAP embedding of cell types of microglia (MGC). (**J**) UMAP embedding of MGC, colored by major brain structures. (**K**) Density scatter plot comparing the averaged accessibility and coefficient of variation across brain structures at each cCRE. Variable cCREs for MGC are defined on the right side of dash line. (**L**) Heat map showing the normalized accessibility of variable cCREs in MGC. (**M**) UMAP embedding of cell types of astrocytes (ASCs). (**N**) UMAP embedding of ASCs, colored by major brain structures. (**O**) Density scatter plot comparing the averaged accessibility and coefficient of variation across brain structures at each cCRE. Variable cCREs for ASCs are defined on the right side of dash line. (**P**) Heat map showing the normalized accessibility of variable cCREs in ASCs. (**Q**) UMAP embedding of non-telencephalon ASCs (ASCNTs). (**R**) UMAP of ASCNTs, colored by major brain structures. (**S**) Normalized chromatin accessibility of 8,790 cell-type-specific cCREs. (**T**) Representative images of transgenic mouse embryos showing LacZ reporter gene expression under the control of the indicated enhancers that overlapped the differential cCRE in **S** (dotted line). Images were downloaded from the VISTA database (https://enhancer.lbl.gov). (**U**) Top enriched known motifs for astrocyte cell-type-specific cCREs.

We found a diverse population of both telencephalic (ASCT) and non-telencephalic astrocytes (ASCNT) in different major brain structures (**Fig. 3M, and N**). We identified three ASCNT cell types from sub-clustering of astrocytes, and one cell population restricted to the cerebellum that was annotated as Bergmann glial cell (ASBGM) (**Fig. 3Q, and R, Table S5**). One cell type (ASCNT-1) was detected mostly in the thalamus, midbrain, and pons, whereas the other two ASCNT cell types were predominantly found in the cortex, hippocampus, and cerebral nuclei (CN). To characterize the dynamic epigenome, we compared the open chromatin landscapes among different cell types using a likelihood ratio test (**Fig. 3S**, **Table S16,** see **Methods**), and identified a total number of 8,790 cCREs that exhibited cell-type-restricted accessibility (range: 100–3,787) (**Fig. 3S**). A human enhancer, specifically accessible in the ASCNT-1 type, was previously validated by mouse transgenics to be active in the midbrain (**Fig. 3T**). We further performed motif analysis for differentially accessible regions in these cell types, finding enrichment of both shared and specific TF binding motifs. For example, we found CCAAT box-binding transcription factor (CTF) NF1 enriched in differential regions identified from both ASCT-1 and ASBGM, whereas TF motifs from nuclear receptors (NRs) and zinc-finger families are specifically enriched in different types (**Fig. 3U, Table S17**).

Medium spiny neurons (MSN) have two primary characteristic types: D1-type and D2-type MSNs defined by the expression of two dopamine receptors DRD1 and DRD2. Although these two sub-classes showed significantly different gene expression patterns and were well separated from each other in transcriptomic and methylation datasets (companion paper, Siletti, et al. 2022 and Wei, et al. 2022), we found that the cell populations of MSNs were well separated by the sub-regions in basal ganglia, rather than D1- and D2-types. (**Fig. S11A-C**). We identified and annotated several cell types as D1-/D2-MSN types based on the chromatin accessibility at gene *DRD1* and *DRD2* loci and sub-regions (**Fig. S11C**). Specifically, we found both D1- and D2-MSN populations from putamen (Pu) (**Fig. S11A-C**). Similarly, D1- and D2-types were also found in the body of caudate (CaB) (**Fig. S11A-C**). We identified one cell type restricted in nucleus accumbens (NAC), although the nuclei also show a distinct pattern of chromatin accessibility at DRD1 and DRD2 loci (**Fig. S11A-C**). We hypothesized that the regional specificity of D1-/D2-MSN types is accompanied by differential chromatin accessibility at the cCREs, which likely underlies cell-specific gene expression patterns. We found a total of 35,203 differential accessible cCREs (**Fig. S11D, Table S18**). Higher chromatin accessibility variation has been found in cell populations from different brain sub-regions, compared to the difference found between D1- and D2-MSN types (**Fig. S11D**). Consistent with above observations, MSN-types were better separated by brain sub-regions, other than separated by D1- and D2-types (**Fig. S11A-C**). The motif analysis suggested the gene expression in these cell types can be regulated by both brain region specific, and D1-/D2- cell-type specific TFs. For example, we found DLX and LHX from the Homeobox TF family were specifically enriched in D1/2-NAC, whereas MEF2 and RORA/G were enriched in both D1- and D2-MSN types from putamen but not in other brain sub-regions (**Fig. S11E, Table S19**). In addition, we found EGR1 and WT1 from zinc-finger TF family were specifically enriched in D2-MSN type but not in D1-MSN types from caudate (**Fig. S11E, Table S19**).

### Conservation of chromatin accessibility at cCREs between mouse and human brain cells

To determine how conserved the gene regulatory landscapes are between human and mouse brains, we compared the human brain cCREs defined in this study with our previously published map of mouse cerebrum cCREs(*40*). We first performed joint clustering of 18 neuronal and glial cell sub-classes (each with >1,000 single nuclei) (**Fig. 4A, Fig. S12**)(*40*). We used multiple molecular features, including “gene activity scores” at homologous genes, chromatin accessibility at homologous cCREs, and transcription factor motif enrichment scores (**Fig. S12A-G,** see **Methods**). We note that clustering based on gene activity scores alone does not align brain cell types between the two species (**Fig. S12A, D, and E**), because of a lack of general conservation of gene expression patterns, as reported previously (*52, 53*). We also identified orthologues of the human cCREs in the mouse genome by performing reciprocal homology searches, and overlapped with cCREs identified in the mouse cerebrum (**Table S20**, **Fig. S12H-I**, see **Methods**). Clustering using the chromatin accessibility at homologous cCREs alone also failed to align corresponding cell types, likely due to significant levels of CRE turnovers (**Fig. S12B, and F**). Instead, we found that clustering based on TF motif enrichment allows for a reasonable alignment of brain sub-classes between species (**Fig. S12C, and G**). We observed that sequence motif enrichment scores were the most conserved molecular features that reliably align similar cell sub-classes between human and mouse brains (**Fig. 4B**, **Fig. S12G**). These analyses suggested that although the open chromatin landscapes were largely divergent between species, the gene regulatory programs of similar cell types may share a similar “vocabulary”, which is the specific combination of transcription factors in each cell lineage.

**Fig. 4:**
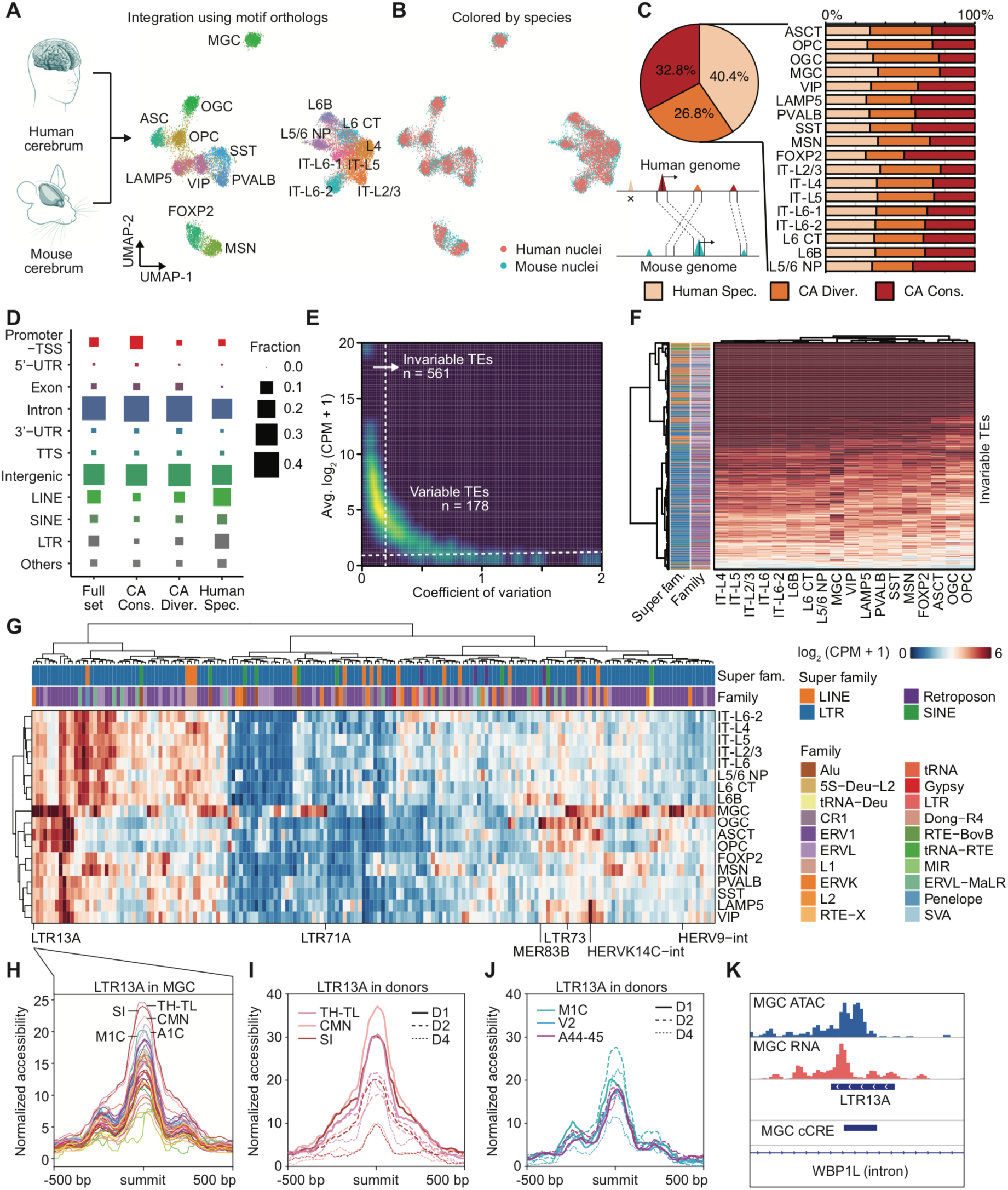
Comparative analyses of chromatin accessibility between human and mouse cerebrum. (**A**) UMAP co-embedding of 18 cell subclasses from both human and mouse cerebrum. (**B**) UMAP co-embedding of single nuclei colored by human and mouse. (**C**) Left, pie chart showing fraction of three categories of cCREs, including human specific, CA-divergent and CA-conserved cCREs. The CA-conserved cCREs are both DNA sequence conserved across species and have open chromatin in orthologous regions. The CA divergent cCREs are sequence conserved to orthologous regions but have not been identified as open chromatin regions in other species. Human specific cCREs are not able to find orthologous regions in the mouse genome. Right, bar plot showing three categories of cCREs in corresponding cell subclasses from human and mouse. (**D**) Dot plot showing fraction of genomic distribution of three categories of cCREs. (**E**) Density scatter plot comparing the averaged accessibility and coefficient of variation across cell subclasses at each transposon elements (TEs). Variable TEs are defined on the upper right side of dash lines, invariable TEs are defined on the upper left of dash lines. (**F**) Normalized accessibility at invariable TEs in different cell subclasses. CPM, counts per million. (**G**) Normalized accessibility at variable TEs in different cell subclasses. (**H**) Average chromatin accessibility of LTR13A in microglia across different brain regions. (**I**) Variable chromatin accessibility of LTR13A across donors in microglia from different brain regions. (**J**) Invariable chromatin accessibility of LTR13A across donors in microglia from different brain regions. (**K**) Representative genomic locus showing chromatin accessibility and expression levels on LTR13A in microglia.

We found that for ∼60% of the human cCREs, mouse genome sequences with high similarity could be identified (more than 50% of bases lifted over to the human genome) (**Fig. 4C, Fig. S12H-I, Table S20**). Among these orthologues’ genome sequences, only half of them (32.8% of total human cCREs) were identified as open chromatin regions in any cell sub-classes from the mouse cerebrum, whereas the other half were not. We thus defined the 32.8% of human cCREs with both DNA sequence similarity and open chromatin conservation as chromatin accessibility (CA) conserved cCREs, and 26.8% of human cCREs with only DNA sequence similarity as chromatin accessibility (CA) divergent cCREs. In addition, we found 40.4% of human cCREs to be without orthologous genome sequences in the mouse genome, and refer to them as human-specific cCREs (**Fig. 4C**, left, **Table S21**). This general pattern was consistent with what has been reported in other cell types between human and mouse(*54*). Next, we further performed the same analyses on the cCREs within the corresponding cell sub-classes from both species. We observed a similar pattern and the proportion of different categories of cCREs was relatively consistent between various cell sub-classes (**Fig. 4C**, right, **Fig. S12J, Table S21**).

We next characterized the genomic distribution of different categories of cCREs. Not surprisingly, we observed that a large proportion of CA-conserved cCREs were located at or near the promoter-TSS regions in the human genome. In addition, the human-specific cCREs were enriched for transposable elements (TE), such as LINEs, SINEs, and LTRs (**Fig. 4D, Table S21**). Most TE families show invariable chromatin accessibility across sub-classes (**Fig. 4E, and F**), and they are dominated by LINE and LTR superfamilies (**Fig. 4F**). Interestingly, 178 TE families display cell-type-specific patterns of chromatin accessibility (**Fig. 4G**). Previous reports suggest that certain transposable elements are active in mammalian brains, and could hypothetically contribute to vulnerability to disease(*55, 56*). Our findings support the hypothesis that distinct TE families might be activated in specific brain cell types. For example, LTRs, including but not limited to LTR13A, LTR71A, and HERV9-int, display chromatin accessibility in microglia but not in other sub-classes of brain cells (**Fig. 4G**). The LTR13A has been reported to act as cellular gene enhancers(*57, 58*). We observed that the LTR13A also shows variable accessibility in microglia populations in different brain regions. For example, we found higher accessibility in brain regions such as posterior parahippocampal gyrus (TH-TL), primary visual cortex (A1C) and primary motor cortex (M1C) from the cortex, substantia innominata and nearby nuclei (SI) and corticomedial nuclear group (CMN) from cerebral nuclei, while lower accessibility in midbrain and cerebellum (**Fig. 4H**). Furthermore, we observed that chromatin accessibilities at LTR13A in microglia from TH-TL, CMN and SI vary considerably among the donors, (**Fig. 4I**), but not in other brain regions (**Fig. 4J**). The chromatin accessibility of LTR13A was associated with the activation indicated by RNA expression signals (**Fig. 4K**), though the biological significance of this observation requires further investigation.

Taking advantage of the integration of multi-modal datasets, we can better delineate the gene regulatory programs that underlie the identity and function of each brain cell type (**Fig. 5A, Fig. S13A-E**). The data integration was anchored on shared genomic features. Here, the RNA expression levels of a common set of genes were profiled from snRNA-seq and Paired-Tag assays. The chromatin accessibility from snATAC-seq at both 2kb upstream and gene body, defined as “gene activity score”, showed a strong correlation with the expression level, which was used for integration(*47*). Given the negative correlation of DNA methylation levels, especially mCG at gene bodies with RNA expression, the expression levels of genes were imputed from both snmC and snm3C-seq data (**Fig. 5A**, see **Methods**). Through the integration of those single cell genomic datasets, we were able to better interrogate the gene regulatory programs in cell types with information from gene expression, open chromatin, histone modification, DNA methylation, and chromosome contact (**Fig. 5A**). We collected single-cell genomic datasets profiled from the human primary cortex (M1C) and middle temporal gyrus (MTG), including 149,891 cells from scRNA-seq, 27,383 cells from Paired-Tag (only from M1C), 55,974 cells from snATAC-seq, 10,604 cells from snmC-seq, and 16,257 cells from snm3C-seq (**Fig. 5B-C**). We performed co-embedding cell clustering analysis on these datasets. We first normalized technical confounders to recover sharper biological heterogeneity using SCTransfrom(*59*). Then, dimensionality reduction (i.e., PCA) was conducted by using a common gene set with Seurat(*60*). Later, the PCA embeddings were iteratively corrected between assays using the Harmony algorithm(*61*). Finally, the cell clusters were identified from the k-nearest neighbor space calculated from the corrected PCA and visualized on UMAP embedding (see **Methods**). The different single-cell assays showed excellent agreement in the same co-embedding space, which indicates the high quality of common variable features and the success of the integration strategy (**Fig. 5C**).

**Fig. 5:**
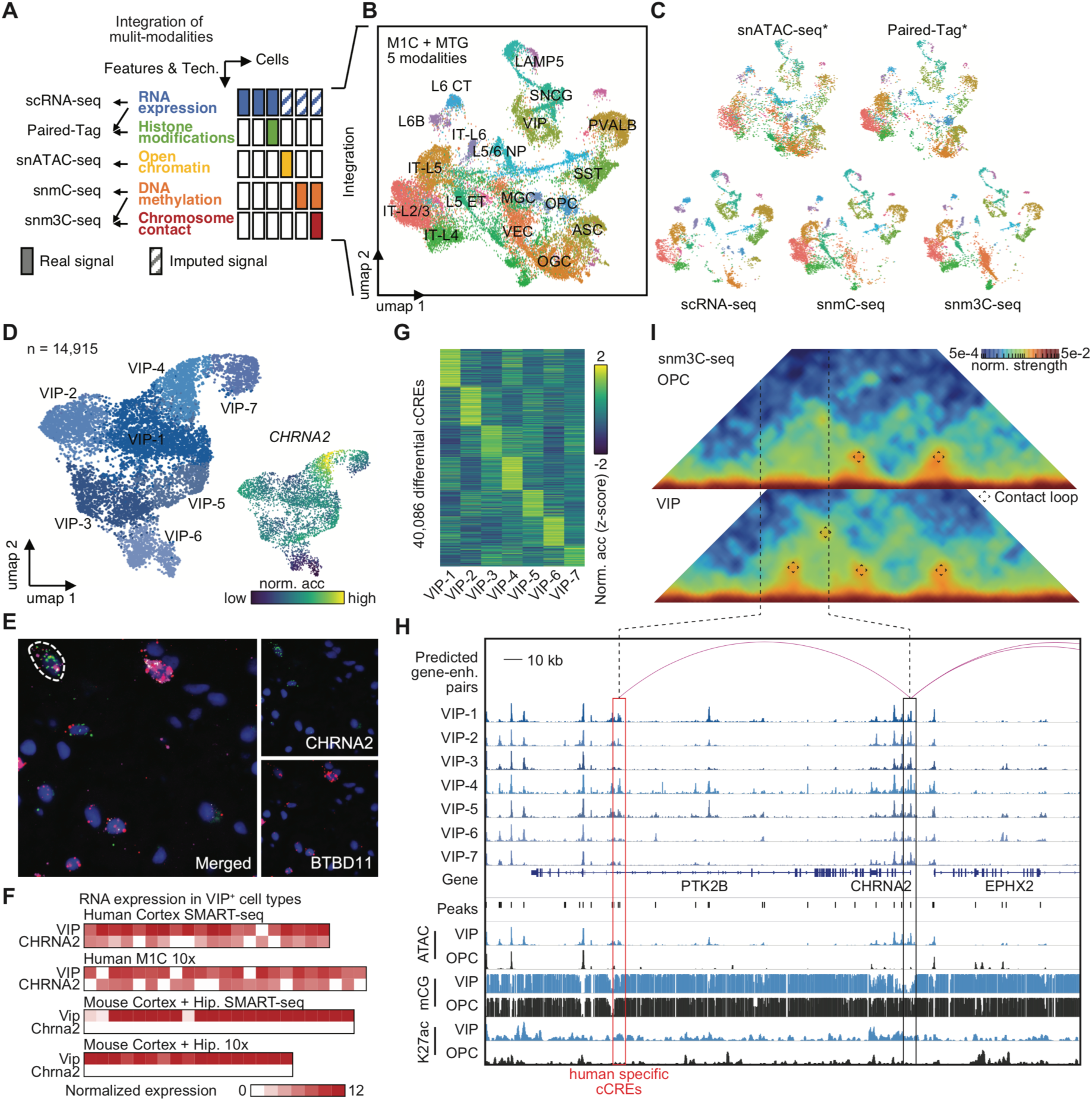
Integration of multi-modal single cell datasets of cortical cells. (**A**) Summary of single cell technologies and multi-model data integration strategies. (B) UMAP embedding and integrative clustering analysis of 18 major cell types. (**C**) Co-embedding of multi-model single cell datasets showing excellent agreement. * for assays using unbiased sampling strategy. (**D**) Left, UMAP embedding of VIP positive (VIP^+^) GABAergic cell types. Upper right, colored by donors. Bottom right, normalized accessibility at gene *CHRNA2*. (**E**) RNAscope validation of CHRNA2-lineage of VIP^+^ GABAergic neurons. (**F**) Expression of gene *VIP* and *CHRNA2* in human VIP+ cell types, and expression of gene *Vip* and *Chrna2* in mouse VIP+ cell types from Allen Cell Types Database: RNA-Seq Data. (**G**) Normalized chromatin accessibility of 40,086 VIP^+^ cell-type-specific cCREs. (**H**) Genome browser track view at the CHRNA2 locus as an example for candidate enhancers predicted from single cell multi-model datasets. Displayed chromatin accessibility profiles from snATAC-seq; DNA methylation signals (mCG) from snm3C-seq; and histone modification signals (H3K27ac) from Paired-Tag for several VIP^+^ neurons and oligodendrocytes precursor cells (OPC). Red Arcs represent the predicted enhancer for gene *CHRNA2*. (**I**) Triangle heat map show chromatin contacts in VIP^+^ neurons and OPC derived from snm3C-seq data at gene *CHRNA2* locus.

We characterized the gene program in VIP-4, one cell type of VIP^+^ neurons, which showed distinct chromatin accessibility at the *CHRNA2* gene locus (**Fig. 5D**). *CHRNA2* encodes a subunit of nicotinic cholinergic receptor, which is involved in fast synaptic transmission. The co-localization of marker gene *BTBD11* for VIP+ neurons and *CHRNA2* from RNAscope(*62*) fluoresence *in situ* hybridization (FISH) experiment first validated the existence of VIP-4 type in the human cortex (**Fig. 5E, Fig. S13F**). We also noticed that the expression of *CHRNA2* was restricted in human VIP+ cell types, but not in any VIP+ cell types identified from mouse brain (**Fig. 5F**) (*9, 63, 64*). We explored whether the human-specific expression of *CHRNA2* was regulated by specific cCREs in the VIP-4 type. We identified a total of 40,086 differential cCREs between 7 VIP^+^ cell types (**Figure 5G, Table S22**). One differential cCRE located downstream of the gene *CHRNA2* showed higher accessibility in VIP-4, than in Oligodendrocyte precursor cells (OPC) (**Fig. 5H**, ATAC tracks for 7 VIP^+^ cell type and aggregated signals for VIP and OPC). This cCRE was also characterized as a human-specific cCRE in VIP+ neurons (**Fig. 4C**, **Table S20**). The specific accessibility of this cCRE and promoter of *CHRNA2* in VIP^+^ neurons were supported by mCG signals from snmC-seq (**Fig. 5H**, mCG tracks). This cCRE was predicted as one putative enhancer that regulates the expression of *CHRNA2* (**Fig. 5H**, red arcs). The potential active function of this CRE was supported by H3K27ac modification from Paired-Tag (**Fig. 5H**, K27ac tracks). We further confirmed the chromatin interactions between this CRE and the promoter of *CHRNA2* (**Fig. 5I**). Taken together, the above data suggested that this human-specific CRE could be an enhancer that regulates the distinct expression of *CHRNA2* in the VIP-4 type from the human brain.

Although the cCREs from human and mouse brains shared considerable evolutionary conservation in DNA sequences, the orthologous elements also displayed a significant amount of divergence in chromatin accessibility in each cell type. We categorized CA-conserved and CA-divergent cCREs in matched cell sub-classes identified from human and mouse brains (**Fig. 4C**). First, we unsurprisingly found ∼40% of the CA-conserved cCREs from every cell sub-class were at TSS proximal genomic regions, compared to ∼10% of CA-divergent cCREs distributed in TSS proximal genomic regions (**Fig. 6A**). Second, we observed that a significantly larger amount of distal CA-conserved cCREs are marked by either H3K27ac or H3K27me3, than CA-divergent cCREs (Wilcoxon signed-rank test, *p*-value < 0.001, **Fig. 6B, and C**, **Table S23,** see **Methods**). Third, the CA-conserved cCREs were largely shared between diverse cell sub-classes, while the CA-divergent cCREs show less overlap between sub-classess and higher cell type-specificity except a few cell sub-classes with close relationship (OPC and OGC, IT neurons, **Fig. 6D**). Fourth, on average, 45% of distal CA-conserved cCREs have been predicted to regulate putative target genes (**Fig. 2F-H**), which is significantly higher than the proportion found in CA-divergent cCREs (**Fig. 6E**). However, the distribution of 1-D genomic distance from putative enhancers to promoters of their putative target genes showed no difference between these two categories of cCREs (**Fig. 6F**).

**Fig. 6:**
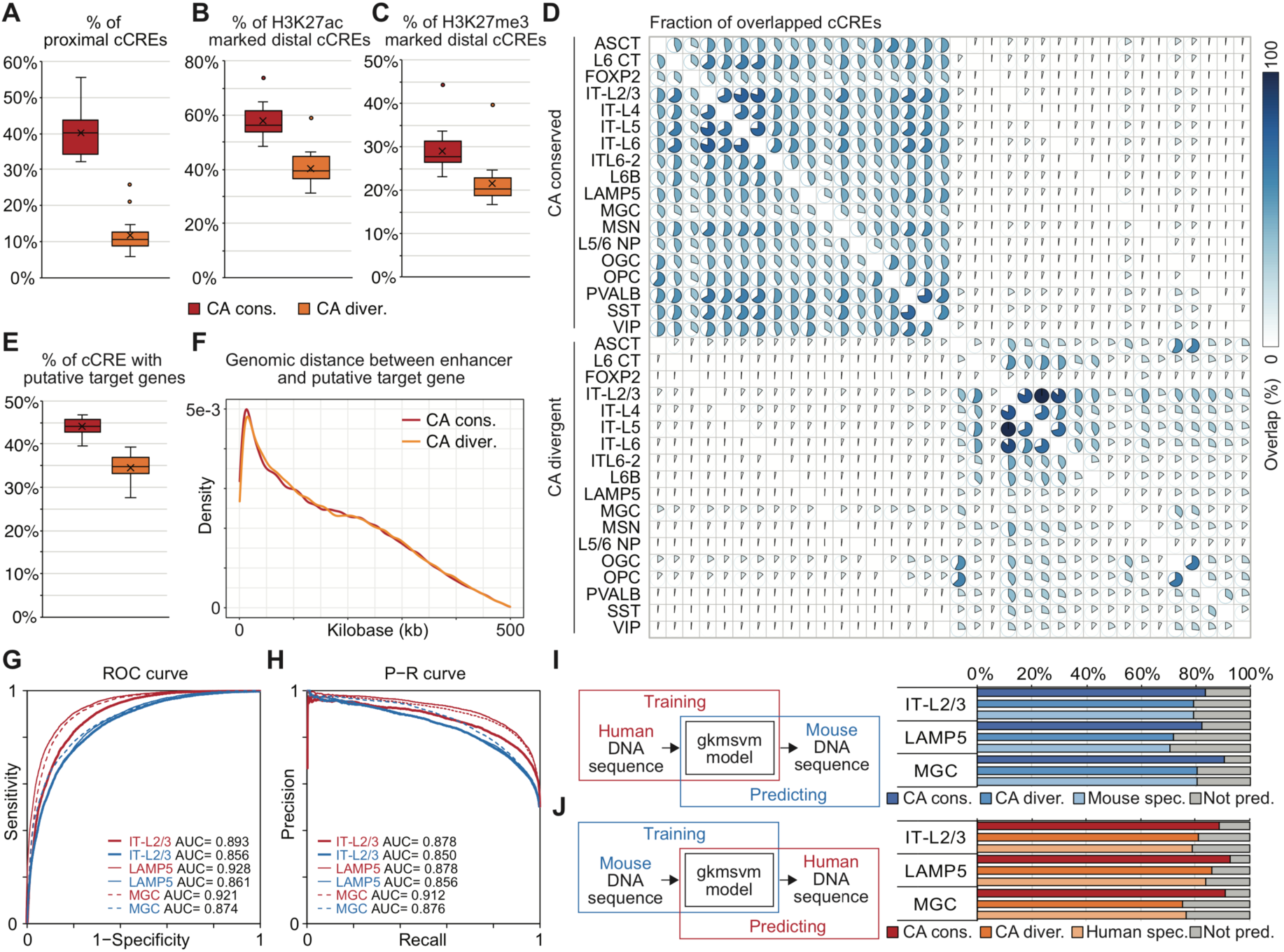
Epigenetic conservation and divergence of human orthologous cCREs. (**A**) The boxplots showing percentage of proximal cCREs in CA-conserved and CA-divergent cCREs from various cell subclasses. (**B**) The boxplots showing percentage of H3K27ac marked distal cCREs in CA-conserved and CA-divergent cCREs from various cell subclasses. (**C**) The boxplots showing percentage of H3K27me3 marked distal cCREs in CA-conserved and CA-divergent cCREs from various cell subclasses. (**D**) Fraction of overlap between CA-conserved and CA-divergent cCREs defined in various cell subclasses. (**E**) The boxplots showing percentage of distal cCREs from various cell subclasses are predicted to regulate putative target genes in CA-conserved and CA-divergent cCREs (Fig. 2). (**F**) The distribution of 1-D genomic distance between epigenetic conserved/divergent enhancers and putative target genes. (**G**) Receiver operating characteristic (ROC) curve and area under curve (AUC) from gkmsvm models trained for representative human and mouse cell types. (**H**) Precision-recall curve (PRC) curve and area under curve (AUC) from gkmsvm models trained for representative human and mouse cell types. (**I**) Prediction for mouse epigenetic conserved, mouse CA-divergent, and mouse specific cCREs from gkmsvm models trained in corresponding human cell subclasses. (**J**) Prediction for human epigenetic conserved, human CA-divergent, and human specific cCREs from gkmsvm models trained in corresponding mouse cell subclasses.

### Sequence changes underlie epigenetic divergence in cCREs in distinct brain cell types

We hypothesize that the epigenetic divergence of cCREs is partly due to evolutionary changes in DNA sequences. To test this hypothesis, we picked IT-L2/3 neurons, LAMP5^+^ interneurons, and MGC as representative cell sub-classes, and trained gapped-kmer SVM classifiers (gkmSVM) (*65*) from the DNA sequences in cCREs (**Fig. 6G-H**, see **Methods**). These models achieved excellent performance (area under receiver operating characteristic curve (AUROC) ranging from 0.856 to 0.928, and area under precision-recall curve (AUPRC) ranging from 0.850 to 0.912) in the prediction of open chromatin regions within the corresponding species (**Fig. 6G-H**). Next, we predicted different categories of mouse cCREs using gkmSVM models trained with human DNA sequence at cCREs in corresponding cell sub-classes (**Fig. 6I**). Importantly, these models also achieved high accuracy (ranging from 0.83 to 0.91) in the prediction of CA-conserved mouse cCREs, and slightly lower accuracy in CA-divergent (ranging from 0.79 to 0.82) and mouse-species cCREs (ranging from 0.71 to 0.80) (**Fig. 6I**). Similarly, the gkmSVM models trained with mouse DNA sequences archived high accuracy in predicting human CA-conserved cCREs (ranging from 0.89 to 0.93), and slightly lower accuracy for human CA-divergent (ranging from 0.75 to 0.86) and human-specific cCREs (ranging from 0.77 to 0.84) (**Fig. 6J**). The human CA-divergent cCREs that failed to be predicted from mouse gkmSVM models have a potential function in regulating genes involved in specific biological processes, including glutamate receptor signaling pathway (GO:0007215), synaptic transmission - GABAergic (GO: 0051932), and various fatty acid elongation (GO: 0019367, GO:0019368, GO:0034625) in IT-L2/3 neurons, LAMP5^+^ interneurons, and MGC, respectively (**Fig. S14, Table S24**). These results suggested that the regulatory divergence is at least in part due to evolution of DNA sequences. However, we cannot exclude a role of non-sequence factors in this process.

### Predicting disease relevant cell types for neuropsychiatric disorders

Genome-wide association studies (GWASs) have identified genetic variants that are associated with many mental diseases and traits (**Table S25**), but >90% of variants are located in non-protein-coding regions of the genome (*4, 5*). Previous studies have shown that non-coding risk variants are enriched in cCREs active in disease relevant cell types (*6–8*). Leveraging the newly annotated cell-type-resolved human brain cCREs, we predict the cell types relevant to the different neuropsychiatric disorders and traits. We performed linkage disequilibrium score regression (LDSC) analysis to determine if the genetic heritability of DNA variants associated with neuropsychiatric traits are significantly enriched within cCREs showing chromatin accessibility in the major brain cell types in the present study (**Table S25**, see **Methods**). We found significant associations between 19 neuropsychiatric disorders and traits (**Table S25, and S26**) with the open chromatin landscapes in one or more cell types we identified (**Fig. 7A**, see **Methods**). In particular, we observed widespread and strong enrichment of genetic variants linked to psychiatric and cognitive traits such as schizophrenia (SCZ), intelligence, bipolar disorder, and educational attainment within accessible cCREs across various neuronal cell types (**Fig. 7A, Table S26**). Other neuropsychiatric traits, such as tobacco use disorder and alcohol usage were associated with specific neuronal cell types in basal ganglia, which were previously implicated in addiction(*66*). Another example is neuroticism, which was restricted and associated with IT neurons from the cortex (**Fig. 7A, Table S26**). In addition, as reported before(*67, 68*), the risk variants from Alzheimer’s disease were significantly enriched in the cCREs found in microglia, but not in other cell types (**Fig. 7A, Table S26**).

**Fig. 7:**
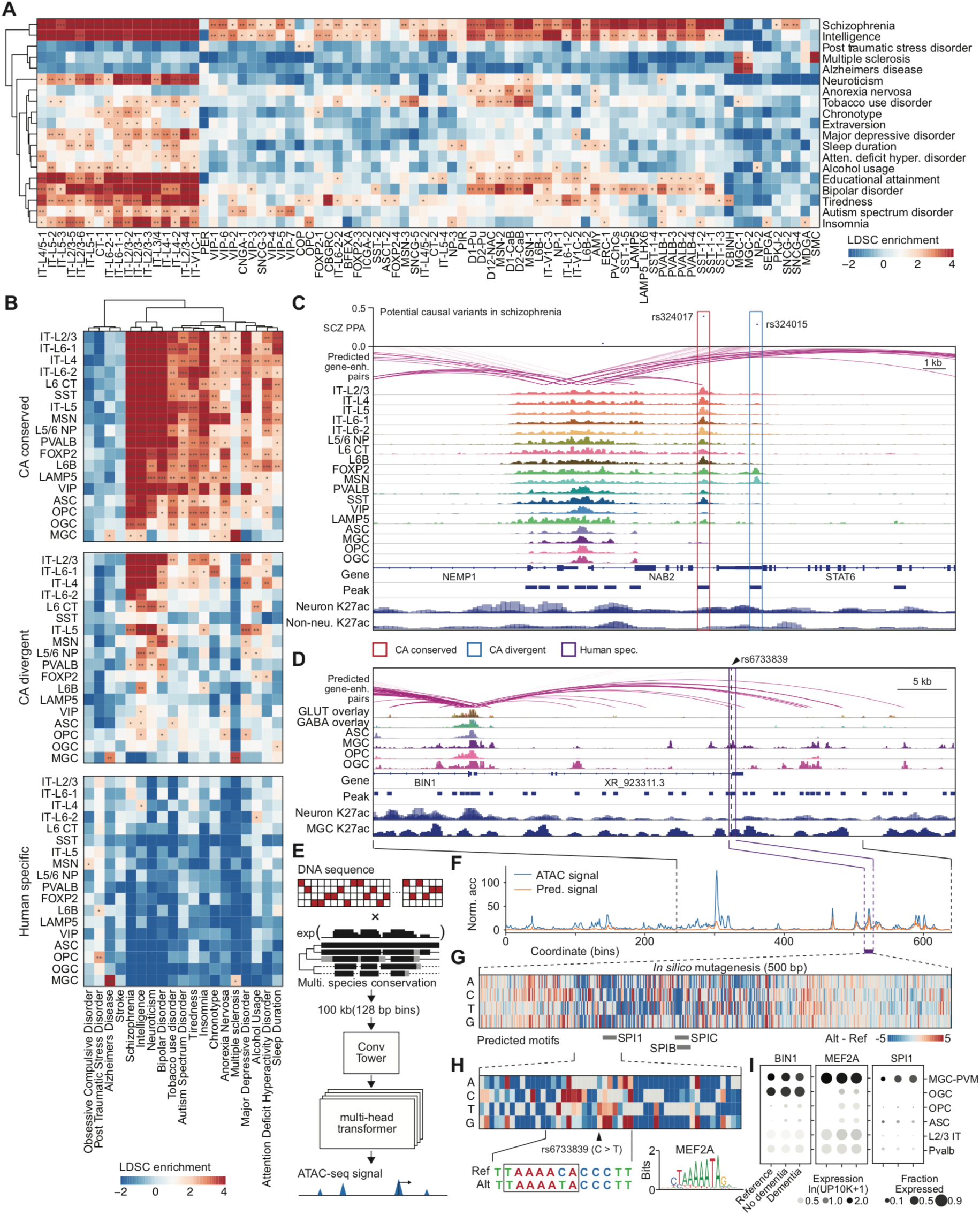
Interpreting noncoding risk variant of neurological disorder and traits. (**A**) Heat map showing enrichment of risk variants associated with neurological disorder and traits from genome wide association studies in human cell type-resolved cCREs. Cell-type specific linkage disequilibrium score regression (LDSC) analysis was performed using GWAS summary statistics. Total cCREs identified independently from each human cell type were used as input for analysis. *P*-values were corrected using the Benjamini Hochberg procedure for multiple tests. FDRs of LDSC coefficient are displayed. *, FDR < 0.05; **, FDR < 0.01; ***, FDR<0.001. Detailed results are reported in **Table S26**. (**B**) Heat map showing enrichment of risk variants associated with mental disorder and traits in three categories of cCREs. Detailed results are reported in **Table S27**. (**C**) Fine mapping and molecular characterization of schizophrenia (SCZ) risk variants in different categories of cCREs from multiple neuronal types. Genome browser tracks (GRCh38) display chromatin accessibility profiles from snATAC-seq; histone modification signals (H3K27ac) from Paired-Tag, and red arcs represent the predicted enhancer for gene *NAB2*. (**D**) Molecular characterization of Alzheimer’s disease (AD) risk variants in human specific enhancer in microglia. Genome browser tracks (GRCh38) display chromatin accessibility profiles from snATAC-seq; histone modification signals (H3K27ac) from Paired-Tag, and red arcs represent the predicted enhancer for gene *BIN1*. PPA: Posterior probability of association. (**E**) Schematic diagram of deep learning model for predicting chromatin accessibly. (**F**) Chromatin accessibility at *BIN1* enhancer loci predicted in human microglia. (**G**) *In silico* nucleotide mutagenesis influenced the prediction of accessibility. Larger signals (in dark red) represent a higher accessibility prediction on altered sequence, and lower signals (in dark blue) represent lower accessibility on altered sequence. (**H**) Higher accessibility predicted on *BIN1* enhancer with risk variant rs6733839 C>T. The risk variant rs6733839 creates a TF motif for MEF2 family. (**I**) Gene expression level of *BIN1*, *MEF2A* and *SPI1* in patients with Alzheimer’s disease collected from Seattle Alzheimer’s Disease Brain Cell Atlas (SEA-AD).

We further provided breakout reports of LDSC analysis by using three categories of cCREs defined above (**Fig. 4C**). We found most significant associations between cell sub-classes and GWAS traits were found in the analysis of epigenetic conserved elements (**Fig. 7B, Table S27**). Fewer significant associations were observed in the analysis of epigenetic divergent elements, and most of GWAS traits showed no significant associations when human-specific elements were used for LDSC (**Fig. 7B**). For example, the risk variants in schizophrenia showed the most significant enrichment in epigenetic conserved elements. We obtained 4,356 likely SCZ causal variants with a posterior probability of association (PPA) score greater than 1% based on Bayesian fine-mapping determined by the Psychiatric Genomics Consortium (https://www.med.unc.edu/pgc/). By overlapping with different categories of elements, we found one schizophrenia-associated locus harboring a CA-conserved element, and another CA-divergent element containing two likely causal variants, rs324017 and rs324015, respectively (**Fig. 7C**). The CA-conserved cCRE, displayed accessibility in multiple neuronal cell types, was predicted to regulate the expression of the SCZ-associated gene *NAB2* (**Fig. 7C**) (*69*). These observations provide new hypotheses regarding the potential functions of noncoding SCZ risk variants.

Interestingly, LDSC analysis using human-specific elements revealed a significant association between Alzheimer’s disease (AD) and microglia (**Fig. 7B, Table S27**). This raises the possibility that the AD-related risk variants could reside in human-specific regulatory elements, and contribute to human-specific gene regulation programs in microglia(*70*). This observation suggests potential limitations of animal models of AD in revealing disease pathology in humans (*71*). One AD risk locus contains multiple microglia-specific cCREs, which cannot find any homologous sequence in the mouse genome (**Fig. 7D**). One of these cCREs harboring AD-risk variants rs6733839 has been predicted to be a microglia-specific enhancer that can regulate the expression of *BIN1* gene, and its function was supported by both H3K27ac modification and previous validation experiment (**Fig. 7D**) (*68*).

### Deep learning models predict the influence of risk variants on gene regulation

To further understand how risk variants contribute to the function of regulatory elements, we used deep learning (DL) models to predict chromatin accessibility from DNA sequences (**Fig. 7E**, **Figure S15A-D**, see **Methods**). The deep learning model architecture was inspired by Enformer (*72*), which adapts attention-based architecture, Transformer, that could better capture syntactic forms (e.g., order and combination of words in a sentence) and outperforms most existing models in natural language processing tasks (*73*). We trained the novel DL model called Epiformer on the normalized pseudo bulk ATAC-seq profiles in human microglia. The resulting model successfully predict the cell type-specific accessibility of cCREs at the *BIN1* locus with a Pearson correlation coefficient of 0.74 (**Fig. 7F**). By contrast, DL models trained using ATAC-seq profiles from other cell types the accessibilities of these cCREs failed to predict the chromatin accessibility profiles from microglia (**Fig. S15F**). To predict the regulatory effects of the risk variant, we further performed *in silico* mutagenesis on the above microglia-specific enhancer near *BIN1* and compared the changes of accessibility predicted from reference and altered DNA sequences. Every nucleotide within this 500 bp enhancer was mutated *in silico*, and the influence on accessibility was measured by assessing the difference between the predicted accessibility for the reference and altered sequences (**Fig. 7G, Table S28,** see **Methods**). Among these *in silico* single nucleotide mutations, most in the flanking regions did not affect the predicted accessibility, while a few nucleotide substitutions significantly increased or decreased the predicted accessibility (**Fig. 7G, Table S28**). The DNA sequence most negatively associated with the predicted accessibility was predicted to contain binding motifs for transcription factors SPI1, SPIB, and SPIC, which bind to the PU-box and play a key role in the differentiation of macrophages, myeloid and B-lymphoid cells(*74, 75*). This result indicates that only specific sequence motifs within the 500bp region are critical for determining the function of CREs. Lastly, the model also predicts that the nucleotide substitutions (C > T / C > G) of AD-risk variants (rs6733839) would increase the accessibility of human microglia specific enhancer (**Fig. 7H**, upper, in dark red, **Table S28**). This risk variant likely creates a new binding motif for MEF2A or other TFs in microglia (**Fig. 7H**, bottom, **Table S28**). Taken together, we hypothesize that this risk variant (rs6733839) will increase the binding and combined effect of TFs SPI1 and MEF2A, which may result in a higher expression level of AD-risk gene *BIN1* in the microglia. These results support a model where the nucleotides within regulatory elements are not equally contributing to function, and capturing the regulatory code of sequence motifs is key to understanding how risk variants influence the potential function of regulatory elements.

## Discussion

Understanding the cellular and molecular basis of brain circuits is one of the grand challenges of the twenty-first century. In-depth knowledge of the transcriptional regulatory program in brain cells would not only improve our understanding of the molecular inner workings of neurons and non-neuronal cells, but could also shed light on the pathogenesis of a spectrum of neuropsychiatric disorders. Here, we report a comprehensive profiling of chromatin accessibility at single-cell resolution in 42 human brain regions. The chromatin accessibility maps of 544,735 cCREs, were probed in >1.1 million nuclei. Taking advantage of our high-resolution brain dissections, we examined the regional specificity in chromatin accessibility of cell types in the human brain and showed that most brain cell types exhibit strong regional specificity. The described cCRE atlas (http://catlas.org) represents a rich resource for the neuroscience community to understand the molecular patterns that underlie the diversification of brain cell types in complementation to other molecular and anatomical data.

Genome-wide association studies (GWAS) have been used to enhance our understanding of polygenic human traits and reveal clinically relevant therapeutic targets for neuropsychiatric disorders. However, our ability to interpret the risk variants has been hampered by a lack of annotation of the non-coding genome. By leveraging both epigenetic conserved, divergent, and human-specific cCRE identified from various human cell types, we prioritize likely causal variants in linkage disequilibrium (LD), link distal cCREs to putative target genes, and predict motifs altered by risk variants using cutting-edge deep learning methods. We revealed hundreds of cell-type trait associations and created a framework to systematically interpret noncoding risk variants.

## Acknowledgments

We would like to thank Dr. Erica Melief, Aimee Schantz, Katie Kern, Amanda Keen, and Lisa Keene of the University of Washington BioRepository and Integrated Neuropathology (BRaIN) Laboratory for outstanding technical and administrative support for tissue collection, preservation, and characterization, and the brain donors and their loved ones, without whom this research would be impossible. We would like to thank the QB3 Macrolab at UC Berkeley for purification of the Tn5 transposase. This publication includes data generated at the UC San Diego IGM Genomics Center utilizing an Illumina NovaSeq 6000 that was purchased with funding from a National Institutes of Health SIG grant (#S10 OD026929). We thank Ethan Armand for discussion and support, Qiurui Zeng for data management and transferring.

## Funding

This work was supported by grants from NIMH U01MH121282 to B.R., M.M.B and J.R.E., UM1 MH130994 to J.R.E., M.M.B. and B.R., NIMH U01MH114812 to E.L and S.L, and in part by the Nancy and Buster Alvord Endowment to C.D.K. The development of deep learning models was sponsored in part by NSF Convergence Accelerator under award OIA-2040727, NIH Bridge2AI Center Program under award U54HG012510, as well as generous gifts from Google, Adobe, and Teradata. Dr. Z.W. is a DDBrown Awardee of the Life Sciences Research Foundation. J.R.E is an investigator of the Howard Hughes Medical Institute.

## Author contributions

Contribution to data generation: Y.E.L., S.P., M.M., N.D.J., C.Z., Z.W., Y.X., A.P.D., N.E., J.O., J.L., L.L., Q.Y., Q.Z., S.E., A.M.Y., J.N., N.D., T.C., N.S., D.H., R.D.H., B. L., C.D.K., A.W. Contribution to data analysis: Y.E.L., Z.W., H.J., O.P., C.K., W.T., K.S., J.S. Contribution to web portal: Y.E.L. Contribution to deep learning model: Y.E.L., Z.W. Contribution to data interpretation: Y.E.L., S.L., E.L., T.B., B.R., M.M.B., J.R.E. Contribution to writing the manuscript: Y.E.L., B.R. All authors edited and approved the manuscript.

## Competing interests

J.R.E is a member of the scientific advisor for Zymo Research and Ionis. B.R. is a co-founder and consultant of Arima Genomics Inc. and co-founder of Epigenome Technologies.

## Data and materials availability

Demultiplexed data can be accessed via the NEMO archive here: https://assets.nemoarchive.org/dat-d6r90fb. Processed data will be available on our web portal CATLAS: http://catlas.org. Additional data are available upon request.

Custom code and scripts used for analysis can be accessed here: https://github.com/yal054/snATACutils and https://github.com/r3fang/SnapATAC.

The deep learning model and pre-trained model can be download from https://github.com/yal054/epiformer.

## Supplementary materials

### Materials and Methods

#### Tissue preparation and nuclei isolation

Tissues from 3 human donors (males aged 29, 42, and 58, R hemisphere) with post-mortem interval <12 hours and RIN score >7.5 were obtained from the Allen Institute. Tissue included 42 regions spanning diverse brain structures, dissected using standard architectonic landmarks and guided by the Allen Brain Human Brain Atlas(*39*). Tissue blocks were separated into 100-150mg chunks in RNase-free glass dishes on dry ice and stored at −80°C until homogenized.

Frozen tissue was homogenized in 4.5 mL Lysis Buffer (0.32 M sucrose, 0.1% Triton X-100, 5 mM CaCl_2_, 3 mM Mg(Ace)_2_, 0.1 mM EDTA pH 8.0, 10 mM Tris-HCl pH 8.0, 1 mM DTT, Roche cOmplete Mini Protease Inhibitor, 24 U/mL Promega RNasin Plus) using glass Dounce Homogenizers. For iodixanol-gradient purification of nuclei, tissue was Dounce homogenized in 5 mL NIMT Buffer (0.25 M sucrose, 0.1% Triton X-100, 25 mM KCl, 5 mM MgCl_2_, 10 mM Tris-Cl pH 8.0, 1 mM DTT, 20 U/mL SUPERase IN, 40 U/mL RNAse OUT), mixed with 50% iodixanol buffer (50% iodixanol (Optiprep), 25 mM KCl, 5 mM MgCl_2_, 20 mM Tris-HCl pH 8.0) to a concentration of 20% iodixanol, and layered onto 25% iodixanol buffer (25% iodixanol (Optiprep), 0.125 M sucrose, 12.5 mM KCl, 2.5 mM MgCl_2_, 5 mM Tris-Cl pH 8.0). Samples were centrifuged at 10,000 x g for 20 min at 4°C in a Beckman Optima-80K Ultracentrifuge with SW-41-Ti swinging bucket rotor. The supernatant was carefully aspirated before nuclei rehydration. All steps were performed on ice using nuclease-free equipment.

#### Single nucleus ATAC-seq using combinatorial indexing

Single nucleus ATAC-seq was performed as described with steps optimized for automation and modifications in permeabilization and sort buffers (*34*). A step-by-step-protocol for library preparation is available here: https://www.protocols.io/view/sequencing-open-chromatin-of-single-cell-nuclei-sn-pjudknw/abstract.

Brain nuclei were pelleted with a swinging bucket centrifuge (500 x g, 5 min, 4°C; 5920R, Eppendorf). Nuclei pellets were resuspended in 1 ml nuclei permeabilization buffer (10mM Tris-HCL (pH 7.5), 10mM NaCl, 3mM MgCl2, 0.1% Tween-20 (Sigma), 0.1% IGEPAL-CA630 (Sigma), 0.01% Digitonin (Promega) and 1% BSA (Proliant 7500804) in molecular biology-grade water) and pelleted again (500 x g, 5 min, 4°C; 5920R, Eppendorf). Nuclei were resuspended in 1 mL high salt tagmentation buffer (36.3 mM Tris-acetate (pH = 7.8), 72.6 mM potassium-acetate, 11 mM Mg-acetate, 17.6% DMF) and counted using a hemocytometer. Concentration was adjusted to 2,000 nuclei/9 µl, and 2,000 nuclei were dispensed into each well of a 96-well plate. For tagmentation, 1 µL barcoded Tn5 transposomes26 were added using a BenchSmart™ 96 (Mettler Toledo, RRID:SCR_018093, Supplementary Table 26), mixed five times, and incubated for 60 min at 37 °C with shaking (500 rpm). To inhibit the Tn5 reaction, 10 µL of 40 mM EDTA was added to each well with a BenchSmart™ 96 (Mettler Toledo, RRID:SCR_018093), mixed 10 times, and the plate was incubated at 37 °C for 15 min with shaking (500 rpm). Next, 10 µL 3 x sort buffer (3 % BSA, 3 mM EDTA in PBS) was added using a BenchSmart™ 96 (Mettler Toledo, RRID:SCR_018093). If processing two or more samples per day, tagmentation was performed with different sets of barcodes in separate 96 well plates. After tagmentation nuclei from individual plates were pooled together, added to a FACS tube and stained with 3 µM Draq7 (Cell Signaling). Using a SH800 (Sony), 20 nuclei per sample were sorted per well into eight 96-well plates (total of 768 wells, 15,360 nuclei per sample) containing 10.5 µL EB (25 pmol primer i7, 25 pmol primer i5, 200 ng BSA (Sigma). Preparation of sort plates and all downstream pipetting steps were performed on a Biomek i7 Automated Workstation (Beckman Coulter, RRID:SCR_018094). After addition of 1 µL 0.2% SDS, samples were incubated at 55 °C for 7 min with shaking (500 rpm). 1 µL 12.5% Triton-X was added to each well to quench the SDS. Next, 12.5 µL NEBNext High-Fidelity 2× PCR Master Mix (NEB) were added and samples were PCR-amplified (72 °C 5 min, 98 °C 30 s, (98 °C 10 s, 63 °C 30 s, 72°C 60 s) × 12 cycles, held at 12 °C). After PCR, all wells were combined. Libraries were purified according to the MinElute PCR Purification Kit manual (Qiagen) using a vacuum manifold (QIAvac 24 plus, Qiagen) and size selection was performed with SPRI Beads (Beckmann Coulter, 0.55x and 1.5x). Libraries were purified one more time with SPRI Beads (Beckmann Coulter, 1.5x). Libraries were quantified using a Qubit fluorimeter (Life technologies, RRID:SCR_018095) and the nucleosomal pattern was verified using a Tapestation (High Sensitivity D1000, Agilent). Libraries generated were sequenced on a NextSeq500 (RRID:SCR_014983, Illumina), HiSeq4000 (RRID:SCR_016386, Illumina) or NovaSeq6000 (RRID:SCR_016387, Illumina) using custom sequencing primers with following read lengths: 50 + 10 + 12 + 50 (Read1 + Index1 + Index2 + Read2). Indexing primers and sequencing primers are in **Table S29**.

#### Processing and alignment of sequencing reads

Paired-end sequencing reads were demultiplexed and the cell index transferred to the read name. Sequencing reads were aligned to mm10 reference genome using bwa(*76*). After alignment, we used the R package ATACseqQC (1.10.2)(*77*) to check for fragment length contribution which is characteristic for ATAC-seq libraries. Next, we combined the sequencing reads to fragments, and for each fragment we performed the following quality control: 1) Keep only fragments quality score MAPQ > 30; 2) Keep only the properly paired fragments with length <1000 bp. 3) PCR duplicates were further removed with SnapTools (https://github.com/r3fang/SnapTools)(78). Reads were sorted based on the cell barcode in the read name.

#### TSS enrichment calculation

Enrichment of ATAC-seq accessibility at TSSs was used to quantify data quality without the need for a defined peak set. The method for calculating enrichment at TSS was adapted from previously described. TSS positions were obtained from the GENCODE database v29 (hg38)(*79*). Briefly, Tn5 corrected insertions (reads aligned to the positive strand were shifted +4 bp and reads aligned to the negative strand were shifted –5 bp) were aggregated ±2,000 bp relative (TSS strand-corrected) to each unique TSS genome-wide. Then this profile was normalized to the mean accessibility ±1,900-2,000 bp from the TSS and smoothed every 11bp. The max of the smoothed profile was taken as the TSS enrichment.

#### Doublet removal

We used a modified version of Scrublet(*80*) to remove potential doublets for every dataset independently. First, we count the reads on gene body and 2 kb upstream of TSS for every single nucleus that passed quality control. Next, cell-by-gene count matrices were used as input, with default parameters, the doublet scores were calculated for both observed nuclei {x_i_} and simulated doublets {y_i_} using Scrublet(*80*). Then, a threshold θ is selected based on the distribution of {y_i_}, and observed nuclei with doublet score larger than θ are predicted as doublets. To determine θ, we fit a two-component mixture distribution by using function normalmixEM from R package mixtools(*81*). The lower component contained majority of embedded doublet types, and the other component contained majority of neo-typic doublets (collision between nuclei from different clusters. We selected the threshold θ where the *p*_1’_ · *pdf* (*x*, µ_1’_, σ_1’_) = *p*_2_ · *pdf*(*x*, µ_2_, σ_2_). This value suggested that the nuclei have same chance of belonging to both classes.

#### Clustering and cluster annotation

We used an iterative clustering strategy using the SnapATAC package(*78*) with modifications as detailed below. For round 1 clustering, we clustered and finally merged single nuclei to three main cell classes: non-neurons, GABAergic neurons, and glutamatergic neurons. For each main cell class, we performed another round of clustering to identify major cell types. Last, for each major cell types, we performed a third round of clustering to find sub-types.

Detailed description for every step is listed below:

1) Nuclei filtering

Nuclei with >=500/1,000 uniquely mapped fragments and TSS (transcription start site) enrichment >5/7 were filtered for individual dataset.

2) Doublet removal:

The potential barcode collisions were also removed for individual datasets.

3) Feature bin selection

First, we calculated a cell-by-bin matrix at 500 kb resolution for every sample independently and subsequently merged the matrices. Second, we converted the cell-by-bin count matrix to a binary matrix. Third, we filtered out any bins overlapping with the ENCODE blacklist (hg38, https://github.com/Boyle-Lab/Blacklist)(82). Fourth, we focused on bins on chromosomes 1-22, X, and Y. Last, we removed the top 5% bins with the highest read coverage from the count matrix.

4) Dimensionality reduction

SnapATAC applies a nonlinear dimensionality reduction method called diffusion maps, which is highly robust to noise and perturbation. However, the computational time of the diffusion maps algorithm scales exponentially with the increase of number of cells. To overcome this limitation, we used adjacency spectral embedding (https://github.com/yixuan/RSpectra) instead of diffusion maps, which can easily scale up to tens of thousands of nuclei without memory issue. We combined the Nyström method (a sampling technique)^68^ to generate the low-dimensional embedding for large-scale dataset.

A Nyström landmark diffusion maps algorithm includes three major steps:

1. sampling: sample a subset of K (K≪N) cells from N total cells as “landmarks”.
2. embedding: compute a diffusion map embedding for K landmarks;
3. extension: project the remaining N-K cells onto the low-dimensional embedding as learned from the landmarks to create a joint embedding space for all cells.

Having more than 1.1 million single nuclei at the beginning, we decided to apply this strategy on level 1 and 2 clustering. 35,000 cells were sampled as landmarks and the remaining query cells were projected onto the diffusion maps embedding of landmarks. Later for the level III clustering, diffusion map embeddings were directly calculated from all nuclei.

5) Batch correction

To correct the potential bias from donors and technical replicates, we applied batch correction on these variables using R package “Harmony”(*61*).

6) Eigenvector selection

To determine the number of eigenvectors of diffusion operator to include for downstream analysis, we generated an “Elbow plot”, to rank all eigenvectors based on the percentage of variance explained by each one. For each round of clustering, we selected the top 10-20 eigenvectors that captured the majority of the variance.

7) Generate K Nearest Neighbor (KNN) Graph

Using the selected significant eigenvectors, we next construct a K Nearest Neighbor (KNN) Graph. Each cell is a node and the k-nearest neighbors of each cell were identified according to the Euclidian distance and edges were drawn between neighbors in the graph. To overcome the limitation on CPU memory and speed on the process, we applied Annoy (Approximate Nearest Neighbors Oh Yeah) instead of the original KNN function(*60*).

8) Graph-based clustering

We applied the Leiden algorithm on the KNN graph using python package leidenalg (https://github.com/vtraag/leidenalg)(83).

9) Optimization on cluster resolution

We tested different ‘resolution_parameter’ parameters (step between 0 and 1 by 0.1) to determine the optimal resolution for different cell populations. For each resolution value, we tested if there was clear separation between nuclei. To do so, we generated a cell-by-cell consensus matrix in which each element represents the fraction of observations two nuclei are part of the same cluster. A perfectly stable matrix would consist entirely of zeros and ones, meaning that two nuclei either cluster together or not in every iteration. The relative stability of the consensus matrices can be used to infer the optimal resolution. To this end, we generated a consensus matrix based on 100 rounds of Leiden clustering with randomized starting seed 5. let M^s^ denote the N × N connectivity matrix resulting from applying Leiden algorithm to the dataset D^s^ with different seeds. The entries of M^s^ are defined as follows:

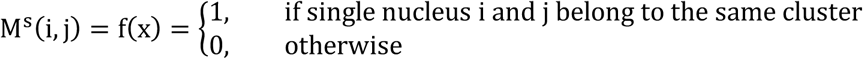

Let I^s^ be the N × N identicator matrix where the (i, j)-th entry is equal to 1 if nucleus i and j are in the same perturbed dataset D^s^, and 0 otherwise. Then, the consensus matrix C is defined as the normalized sum of all connectivity matrices of all the perturbed D^s^.

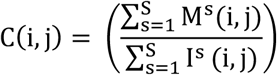

The entry (i, j) in the consensus matrix is the number of times single nucleus i and j were clustered together divided by the total number of times they were selected together. The matrix is symmetric, and each element is defined within the range [0,1]. We examined the cumulative distribution function (CDF) curve and calculated proportion of ambiguous clustering (PAC) score to quantify stability at each resolution. The resolution with a local minimum of the PAC scores denotes the parameters for the optimal clusters. In the case these were multiple local minimal PACs, we picked the one with higher resolution. Another measurement is dispersion coefficient (DC), which reflects the dispersion (ranges from 0 to 1) of the consensus matrix M from the value 0.5. The closer to 1 is the DC, the more perfect is consensus matrix, and thus the more stable is the clustering. In a perfect consensus matrix, all entries are 0 or 1, meaning that all connectivity matrices are identical. The DC is defined as:

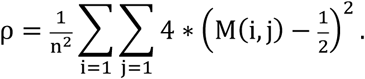

Finally, for every cluster, we tested whether we could identify differential features compared to all other nuclei (background) and the nearest nuclei (local background) using the function ‘findDAR’.

9) Visualization

For visualization, we applied Uniform Manifold Approximation and Projection (UMAP)(*84*).

#### Dendrogram construction for mouse brain cell types

First, we calculated for cCRE the median accessibility per cluster and used this value as cluster centroid. Next, we calculated the coefficient of variation (CV) for the cluster centroid of each element across major cell types. Finally, we only kept variable elements with CV larger than 10% quantile and smaller than 90% quantile for dendrogram construction.

We used the set of variable features defined above to calculate a correlation-based distance matrix. Next, we performed linkage hierarchical clustering using the R package pvclust (v.2.0) (*85*)with parameters method.dist=“cor” and method.hclust=“ward.D2”. The confidence for each branch of the tree was estimated by the bootstrap resampling approach.

#### Regional specificity of cell types

The specificity score is defined as Jensen–Shannon divergence, which measures the similarity between two probability distributions. For each cell sub-class, the contribution of different brain regions is first calculated. Then, we compared this distribution with the contribution of brain regions calculated from all sampled cells. We used function ‘JSD’ from R package philentropy for this analysis(*86*).

#### Identification of reproducible peak sets in each cell cluster

We performed peak calling according to the ENCODE ATAC-seq pipeline (https://www.encodeproject.org/atac-seq/). For every cell cluster, we combined all properly paired reads to generate a pseudo-bulk ATAC-seq dataset for individual biological replicates. In addition, we generated two pseudo-replicates which comprise half of the reads from each biological replicate. We called peak for each of the four datasets and a pool of both replicates independently. Peak calling was performed on the Tn5-corrected single-base insertions using the MACS2(*87*) with these parameters: --shift -75 --extsize 150 --nomodel --call-summits --SPMR -q 0.01. Finally, we extended peak summits by 250 bp on either side to a final width of 501 bp for merging and downstream analysis. To generate a list of reproducible peaks, we kept peaks that 1) were detected in the pooled dataset and overlapped ≥ 50% of peak length with a peak in both individual replicates or 2) were detected in the pooled dataset and overlapped ≥ 50% of peak length with a peak in both pseudo-replicates.

We found that when cell population varied in read depth or number of nuclei, the MACS2 score varied proportionally due to the nature of the Poisson distribution test in MACS2(*87*). Ideally, we should perform a reads-in-peaks normalization, but in practice, this type of normalization is not possible because we don’t know how many peaks we will get. To account for differences in performance of MACS2(*87*) based on read depth and/or number of nuclei in individual clusters, we converted MACS2 peak scores (-log10(q-value)) to “score per million”^72^. We filtered reproducible peaks by choosing a “score per million” cut-off of 2 was used to filter reproducible peaks.

We only kept reproducible peaks on chromosome 1-19 and both sex chromosomes, and filtered ENCODE blacklist (hg38, https://github.com/Boyle-Lab/Blacklist)(82). A union peak list for the whole dataset was obtained by merging peak sets from all cell clusters using BEDtools(*88*).

#### Computing chromatin accessibility scores

Accessibility of cCREs in individual clusters was quantified by counting the fragments in individual clusters normalized by read depth (counts per million: CPM). For each gene, we summed up the counts within the gene body + 2kb upstream to calculate “gene activity score (GAS)” GAS was used for integrative analysis with other single cell modalities. For better visualization, we smoothed GAS to 50 nearest neighbour nuclei using Markov Affinity-based Graph Imputation of Cells (MAGIC)(*89*).

#### Integrative analysis of single nucleus ATAC-seq and single cell datasets

For integrative analysis, we first filtered brain regions that matched samples profiled in this study (same brain dissections were used in companion papers). Second, we manually subset cell types into two groups, neurons and non-neurons, by checking both signal on marker genes and taxonomy labels from scRNA-seq, snmC-seq, and snm3C-seq. To directly compare our single nucleus chromatin accessibility derived cell clusters with the single cell transcriptomics defined taxonomy of the human brain, we first used the snATAC-seq data to impute RNA expression levels according to the chromatin accessibility of gene promoter and gene body as described previously. For single cell methylome, snmC-seq and snm3C-seq, the signal on gene body was negatively correlated with the gene expression. We imputed the RNA expression levels from methylation signals by subtracting it from 1.

Next, we used a 3-step method analogous to Seurat v3 to project three modalities onto the same space: 1) Using canonical correlation analysis (CCA) to capture the shared variance across cells between datasets; 2) finding anchors as 50 mutual nearest neighbors (MNN) between the two datasets; 3) pulling the three modalities into the same space. Detailed description can be found in companion paper (Wei, et al. 2022). To quantify the similarity between cell clusters from two modalities, we calculated an overlapping score as the sum of the minimum proportion of cells/nuclei in each cluster that overlapped within each co-embedding cluster(*90*). Cluster overlaps varied from 0 to 1 and were visualized as a heat map with snATAC-seq clusters in rows and scRNA-seq clusters in columns. We found strong correspondence between the two modalities, for example scRNA-seq and snATAC-seq, which was evidenced by co-embedding of both transcriptomic (T-type) and chromatin accessibility (A-type) cells in the same joint clusters, as well as co-embed with methylation (M-type) cells (**Fig. S6**). For this analysis, we examined neurons and non-neuronal cell classes separately, due to the CG methylation was more informative for non-neurons comparing to CH methylation.

#### Identification of *cis* regulatory modules

We used Nonnegative Matrix Factorization (NMF)(*91*) to group cCREs into *cis* regulatory modules based on their relative accessibility across major clusters. We adapted NMF (Python package: sklearn(*92*) to decompose the cell-by-cCRE matrix V (N×M, N rows: cCRE, M columns: cell clusters) into a coefficient matrix H (R×M, R rows: number of modules) and a basis matrix W (N×R), with a given rank R:

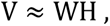

The basis matrix defines module related accessible cCREs, and the coefficient matrix defines the cell cluster components and their weights in each module. The key issue to decompose the occupancy profile matrix was to find a reasonable value for the rank R (i.e., the number of modules). Several criteria have been proposed to decide whether a given rank R decomposes the occupancy profile matrix into meaningful clusters. Here we applied two measurements “Sparseness”(*93*) and “Entropy”(*94*) to evaluate the clustering result. Average values were calculated from 100 times for NMF runs at each given rank with random seed, which will ensure the measurements are stable.

Next, we used the coefficient matrix to associate modules with distinct cell clusters. In the coefficient matrix, each row represents a module and each column represents a cell cluster. The values in the matrix indicate the weights of clusters in their corresponding module. The coefficient matrix was then scaled by column (cluster) from 0 to 1. Subsequently, we used a coefficient > 0.1 (∼95th percentile of the whole matrix) as threshold to associate a cluster with a module.

In addition, we associated each module with accessible elements using the basis matrix. For each element and each module, we derived a basis coefficient score, which represents the accessible signal contributed by all cluster in the defined module. In addition, we also implemented and calculated a basis-specificity score called “feature score” for each accessible element using the “kim” method(*94*). The feature score ranges from 0 to 1. A high feature score means that a distinct element is specifically associated with a specific module. Only features that fulfill both following criteria were retained as module specific elements:

1. feature score greater than median + 2 standard deviations;
2. the maximum contribution to a basis component is greater than the median of all contributions (i.e. of all elements of W).

#### Identification of differentially accessible regions and definition of specificity score

To identify cCRE that differentially accessible in either in cell cluster or brain region, we constructed a logistic regression model predicting cluster/region membership based on each cCRE individually and compares this to a null model with a likelihood ratio test. We used two functions “*fit_models*” and “*compare_models*” in R package Monocle3 (v0.2.2)(*95*) to perform the differential test. We designed the full model as

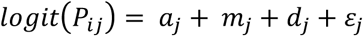

and a reduced mode as

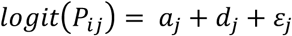

Where *P_ij_* represents the probability of i^th^ site is accessible in the j^th^ cell, s is the log_10_ transformed total number of sites observed as accessible for the jth cell, u is membership of the j^th^ cell in either cluster or region being tested, *d* is the donor label for j^th^ cell and *ε* is an error term.

For each set of testing, between cell type or between regions in every cell sub-class, we only kept cCREs that overlapped with peaks identified in corresponding cell types. A likelihood ratio test is then used to determine if the full model (including cell cluster or region membership) provided a significantly better fit of the data than reduced model. After correcting *p*-value using Benjamini-Hochberg method, we set an FDR cutoff as 0.001 to filter out significant differential cCREs.

Log_2_ fold change is used for two-group comparison, for multiple groups, we calculated a Jensen Shannon divergence-based specificity score described in previous study,^22^ in order to better assign differential cCREs to cell type or brain region. The fraction of accessibility of each cluster + was first calculated for each i^th^ site. We normalized these scores by multiplying by corresponding scaling factors, which are considering different overall complexity across groups. To do so, median number of sites accessible w in individual cells for each cluster was calculated and followed with log_10_-transforming. Then, we took the ratio of the average of w (across all clusters) over the median accessibility in each cluster as scaling factor. These corrected fraction of accessibility for each cCRE was then converted to probability by scaling by groups. Then, we calculated Jensen Shannon divergence between two probability distributions. For example, the probability distribution for the 1^st^ cCRE as *1, to test whether this cCRE is specific in group1, we assumed another probability distribution:

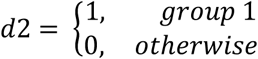

Function “*JSD”* in R package “philentropy” was used to calculate Jensen Shannon divergence between these two probability distributions, and Jensen-Shannon-based specificity scores (JSS) was defined as:

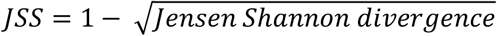

For each group, we calculated JSS for every cCRE. To find a reasonable cut-off for determining restricted or general cCREs, we consider JSS from all cCREs that are not identified as differential accessible (from likelihood ratio test) as a background distribution, and JSS from cCREs that passed our likelihood ratio test threshold and had positive values to be true positives. We set an empirical FDR cut-off at where type I error no more than 5%.

Finally, the differential cCREs could be aligned to multiple subtype or brain regions based on JSS, we named the one can be assigned to only one type or region as region-specific or cell-type-specific cCREs.

#### Predicting enhancer-promoter interactions

First, co-accessible regions are identified for all open regions in each cell cluster (randomly selected 200 nuclei, and used all nuclei for cell cluster with <200 nuclei) separately, using Cicero(*47*) with following parameters: aggregation k = 10, window size = 500 kb, distance constraint = 250 kb. In order to find an optimal co-accessibility threshold for each cluster, we generated a random shuffled cCRE-by-cell matrix as background and identified co-accessible regions from this shuffled matrix. We fitted the distribution of co-accessibility scores from random shuffled background into a normal distribution model by using R package fitdistrplus(*96*). Next, we tested every co-accessibility pairs and set the cut-off at co-accessibility score with an empirically defined significance threshold of FDR<0.01.

The cCREs outside ± 2 kb of transcriptional start sites (TSS) in GENCODE hg38 (v29)(*97*) were considered distal. Next, we assigned co-accessibility pairs to three groups: proximal-to-proximal, distal-to-distal, and distal-to-proximal. In this study, we focus only on distal-to-proximal pairs. We calculated Pearson’s correlation coefficient (PCC) between gene expression and cCRE accessibility across matched T-type and A-type clusters to examine the relationship between co-accessibility pairs. To do so, we first aggregated all nuclei/cells from scRNA-seq and snATAC-seq for every joint cluster to calculate accessibility scores (log_2_ CPM) and relative expression levels (log_2_ normalized UMI). Then, PCC was calculated for every gene-cCRE pair within a 1 Mbp window centered on the TSS for every gene. We also generated a set of background pairs by randomly selecting regions from different chromosomes and shuffling of cluster labels. Finally, we fit a normal distribution model and defined a cut-off at PCC score with empirically defined significance threshold of FDR<0.01, in order to select significant positively correlated cCRE-gene pairs.

#### RNAscope fluorescence *in situ* hybridization

1. Tissue preparation: All tissues were cryostat sectioned at 14 µm onto SuperFrost Plus charged slides and allowed to adhere to the slide. The slides were then stored at −80 C until further use.
2. RNAscope in situ hybridization multiplex version 2 was performed as per the protocol established by Advanced Cell Diagnostics (ACD) with minor modifications. In brief, slides were fixed using cold (4°C) 4% PFA for 15 minutes. The samples were then dehydrated in 50% ethanol (5 min), 70% ethanol (5 min) and 100% ethanol (5 min, twice) at room temperature. The slides were air dried for 10 min, after which boundaries were drawn around each section using a hydrophobic pen (ImmEdge PAP pen; Vector Labs). After the hydrophobic boundaries dried, sections were incubated with hydrogen peroxide (ACD, ∼5 drops per section) for 10 MIN at RT. Sections were then washed twice with PBS for 2 MIN at RT and, subsequently, incubated with 4-5 drops of RNAscope® Protease IV (15 min) at RT. Slides were washed twice in 1X phosphate buffered saline (PBS, pH 7.4) at room temperature (2 min each). Each slide was then placed in a prewarmed humidity control tray (ACD) containing dampened filter paper, and a mixture of Channel probes (50:1:1 dilution, as per ACD instructions) was pipetted onto each section until fully covered. The humidity control tray was placed in a HybEZ oven (ACD) for 2 hours at 40°C. Following probe incubation, the slides were washed twice in 1X RNAscope wash buffer and stored overnight in 5X SSC (Sigma-Aldrich) at RT. The following day slides were returned to the oven for 30 minutes after submersion in AMP-1 reagent (∼5 drops). Two washes and amplification were repeated using AMP reagents with a 30-min, 30-min, and 15-min incubation period, respectively. HRP-C1 (ACD, ∼5 drops) was added to the slide and incubated in the HybEZ oven at 40°C for 15 min. Two washes with 1X wash buffer (2 min each) were performed and 150 µL pre-assigned Opal dye added (Opal Dyes; Akoya Biosciences, Opal Dye diluted 1:1500 in TSA buffer). The slides were incubated in the oven for 30 minutes before being washed twice in 1X wash buffer. Slides were then covered in HRP blocker (ACD, ∼5 drops) and incubated for 15 minutes before being washed twice. Slides were then washed two times in 0.1M phosphate buffer (PBS, pH=7.4) and incubated with DAPI (ACD, ∼5 drops) for 30 seconds before being washed in PBS, air dried, and cover-slipped with Prolong Gold Antifade mounting medium. Slides were stored at 4°C until imaging.
3. Image quantification: Fluorescence signals were acquired using a a Zeiss AiryScan confocal laser scanning microscope equipped with a Plan-APOCHROMAT 63× oil-immersion lens (N/A 1.4). Six fields of view (three per layer) of one tissue section were analyzed using Imaris (Bitplane). Prior to quantification, regions containing lipofuscin signals (defined by overlapping non-specific signals appearing in both channels) were subtracted from each image. Then, positive-stained nuclear signals, defined by automated surface detection function, were segmented, and classified into two groups depending on whether they expressed Gad2. Finally, Chrna2 expression within those two groups was quantified.

#### Paired-Tag experimental procedure

Paired-Tag experiment was performed as previously described(*98*) with minor modifications detailed below. First, antibodies against H3K27ac (Abcam, ab4729) or H3K27me3 (Abcam, ab195477) were pre-incubated with protein A-Tn5 fusion protein, and then incubated with permeabilized nuclei overnight, to target the binding of protein A-fused Tn5 to chromatin. Tagmentation reaction and reverse transcription (RT) were then sequentially performed. Reactions were carried out in 12 different wells, each well containing roughly 2M nuclei and a well-specific DNA barcode included in the Tn5 transposase adaptors and RT primers, to label different samples or replicates. Next, a ligation-based combinatorial barcoding strategy was used to introduce the 2^nd^ and 3^rd^ rounds of DNA barcodes to the nuclei, by sequentially attaching well-specific DNA barcodes to the 5’-end of both chromatin DNA fragments and cDNA from RT in 4 x 96-well plates in each round. Finally, the barcoded nuclei were divided into sub-libraries of 25,000 nuclei each and lysed with Proteinase K (NEB, #P8102). The chromatin DNA and cDNA were subsequently purified as a whole with SPRI beads (Beckman Coulter), pre-amplified, and splitted into two parts for RNA and DNA libraries. For the RNA part, libraries were constructed using the Nextera XT DNA Library Preparation Kit (Illumina, FC-131-1024). For the DNA part, an adaptor containing Truseq i5 sequencing primer binding site was ligated to the library fragments, followed by DNA cleanup using SPRI beads and indexing PCR. Final libraries were quantified using qPCR, and sequenced on a NovaSeq 6000 platform (Illumina).

#### Paired-Tag data processing

A detailed, step-by-step Paired-Tag data processing pipeline can be found at: https://github.com/cxzhu/Paired-Tag.

1. Preprocessing: Cellular barcodes from the sequencing reads are first extracted by matching the linker sequences adjacent to the cellular barcodes, which are then mapped to the cellular barcodes reference using bowtie(*99*), with reads with more than 1 nt mismatch discarded. The adaptor sequences are trimmed from 3’ of DNA and RNA libraries, with Poly-dT and random hexamer sequences further trimmed from 3’ of RNA libraries. The low-quality reads (L = 30, Q = 30) are excluded from further analysis. DNA reads are mapped to hg38 using bowtie2(*100*) with default parameters. Reads with MAPQ<10 are removed. PCR duplicates are also removed.
2. Cell clustering: RNA alignment files are converted to a matrix with cells as columns and genes as rows. Cells with less than 200 features in both DNA and RNA matrices are removed. The clustering of single-cells based on RNA-profiles was performed with Seurat package(*60*). Briefly, cell-to-gene counts are normalized, and the variable genes are selected for dimension reduction by PCA. Batch effects are corrected with Harmony(*61*), and single cell gene expression profiles are visualized with UMAP and clustered with Louvain algorithm(*83*).
3. Cell-type-specific histone modification maps: After clustering, the DNA component of Paired-Tag data corresponding to each histone mark in each cell cluster is aggregated to generate profiles of histone modification for the cluster. The aggregated aligned reads were then used to generate genome browser tracks to reveal the landscape of each histone modification in each cell type as defined by the RNA modality from the Paired-Tag data.
4. To annotate each cCRE, we counted sequencing reads on cCREs captured from Paired-Tag experiments on two histone modifications, and fit a Poisson distribution to filter out cCREs that significantly (FDR<0.05) marked by H3K27ac and H3K27me3.

#### Integration analyses of chromatin accessibility between human and mouse brain cell types

To better compare the open chromatin landscape within a similar cellular context, I extracted 18 major neuronal and glial cell types from the mouse cerebrum (*40*) for alignment with human snATAC-seq from this study. We first calculated chromatin accessibility at orthologous genes and cCREs, and enrichment of orthologoues motifs from pseudo-bulk profiles in each cell sub-classes. Then, we preformed multidimensional scaling analyses and project cell classes on two diamentional spaces.

To integrate at single cell level, we randomly selected 1000 nuclei from each cell sub-class in two species. Similarly, we first calculated fragment counts on orthologous genes and cCREs, and variability of orthologous motifs using chromVAR(*101*). For the integration analyses using orthologous genes and motifs as features, we generated Seurat objects in R. Next, variable features were identified and used for identifying anchors between cells from two species. Finally, to visualize all the cells together, we co-embedded the cell profiles from two species in the same low dimensional space. For the integration analysis using orthologous cCREs, we performed by using the SnapATAC package(*78*) as described above. Of note, we did not perform any correction for potential bias from species.

#### Deep learning model Epiformer

The deep learning model takes both one-hot-encoded DNA sequences (A = [1, 0, 0, 0], C = [0, 1, 0, 0], G = [0, 0, 1, 0], T = [0, 0, 0, 1]) and conservation scores (phastCons, range from 0 to 1) calculated from multiple species alignment as inputs. We first divide the human genome (hg38) into segments in length of 98,304 bp. Any genomic segments overlapped with human ENCODE “blacklist” regions, or contain ‘N’ are discarded. The remaining one-hot-encoded DNA sequences are then multiplied by the exponential transformed conservation score:

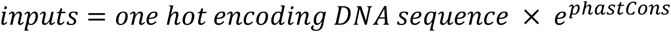

The ATAC-seq signals on the corresponding genomic segments are treated as the predicting targets. We first generate and normalize the genomic signal tracks by using deepTools(*102*) with parameter “--normalizeUsing RPKM”. Then, each segment is cut into 768 genomic bins in size of 128-bp, the mean value of ATAC signal across 128 bp is taken and assigned to that bin. To avoid potential bias from outlier signals, we cutoff signal values higher than 128, so that the bin signal is in the range of 0 to 128.

The model architecture is inspired by Enformer(*72*) with some modifications (see Supplementary **Fig. S15**). It is written based on one open-source machine learning framework, PyTorch, and contains three major parts: [1] 6 residual convolutional blocks with max pooling (Conv Tower). The kernel size (4, 15) is used for the 1^st^ residual convolutional block, and kernel size (4, 5) is used for the rest of residual convolutional block with padding, in order to extract informative sequence features at lower and higher resolution, respectively; The batch normalization and Gaussian Error Linear Unit (GELU) activation function are inserted between each convolutional layer. The convolutional blocks with pooling reduce the spatial dimension from 98,304 bp to 768 so that each sequence position vector represents a bin of 128 bp. [2] Transformer blocks including 4 multi-head attention layers is followed by the Conv Tower, to capture the long-range combinations and orders of sequence features. To inject positional information, we add relative positional encodings(*103*). Dropout rate of 0.4 is used for attention layer to avoid potential overfitting. The SwiGLU(*104*), a GELU variant, is used as activation function. [3] A pointwise convolution block is included to aggregate the weights from the last transformer layer and eventually output the predicted signals. The GELU and SwiGLU activation functions are used with dropout rate of 0.1.

To train the model, we randomly selected 5,000, 10,000 and full set of inputs together with its targets. We found that the model performance increase with the size of training set (**Fig. S15**). To balance the training time and performance, we used 10,000 inputs together with its target for training. Among 80% of inputs are used as training set, and the rest of 20% are used as validation set. To avoid overfitting problem, we introduce data augmentation during training by randomly shifting the input sequence by up to 3 bp and reverse-complementing the input sequence while reversing the targets every other epoch. We used the Adam optimizer from PyTorch with a learning rate of 0.0001 and default settings for other hyperparameters. The model was trained on 4 GPUs (NVIDIA GeForce RTX 3090 24GB). To overcome the limitation in GPU memory, we finally used a gradient accumulation approach to save the gradients and update network weights every 8 batches, to achieve an effective batch size of 64 (16 per GPU). We implemented a cosine warmup learning rate scheduler by linearly increase the learning rate from 0 to target value in the first 10 epoches, and then decrease the learning rate to 0 by multiplying the cosine function in the next 90 epoches. The negative log likelihood loss with Poisson distribution of target (PoissonNLLLoss) is used for evaluating how well the model predict the targets. A cropping layer is introduced to trims 64 positions on each side to avoid computing the loss on the far ends because these regions are disadvantaged because they can observe regulatory elements only on one side (toward the sequence center) and not the other (the region beyond the sequence boundaries).

To evaluate the model, we test the model performance during the training on validation set, by calculating both loss and Pearson Correlation Coefficient (PCC) between predicted and target values across every input in batch. The model checkpoints with either lowest loss or highest PCC have been saved. We finally picked with model with lowest loss for downstream analyses. To evaluate the influence of risk variants on predicted chromatin accessibitly, we performed *In silico* mutagenesis by altering every nucleotide within one cCRE. The predicted signals were calculated for both reference and alterative sequence for every output genomic position/bin. We then took the sum of predicted signals on output positions/bins that overlapped with cCRE, and measure the difference by substracting the sum value on reference sequence from the sum value on alternative sequences.

#### Motif enrichment

We performed both *de novo* and known motif enrichment analysis using Homer (v4.11)(*105*). For cCREs in the consensus list, we scanned a region of ± 250 bp around the center of the element. Randomly selected background regions are used for motif discovery.

#### Gene ontology enrichment

We perform gene ontology enrichment analysis using R package Enrichr(*106*). Gene set library “GO_Biological_Process_2018” was used with default parameters. The combined score is defined as the p-value computed using the Fisher exact test multiplied with the z-score of the deviation from the expected rank.

#### GWAS enrichment

To enable comparison to GWAS of human phenotypes, we used liftOver with default setting “-minMatch=0.95” to convert accessible elements from hg38 to hg19 genomic coordinates(*107*). Next, we reciprocal lifted the elements back to mm10 and only kept the regions that mapped to original loci. We further removed converted regions with length > 1kb.

We obtained GWAS summary statistics for quantitative traits related to neurological disease and control traits (**Table S25**): Alcohol Usage(*108*), Alzheimer’s Disease(*109*), Anorexia Nervosa(*110*), Attention Deficit Hyperactivity Disorder(*111*), Autism Spectrum Disorder(*112*), Bipolar Disorder(*113*), Chronotype(*114*), Educational Attainment(*115*), Extraversion(*116*), Insomnia(*117*), Intelligence(*118*), Major Depressive Disorder(*119*), Multiple sclerosis(*120*), Neuroticism(*121*), Posttraumatic Stress Disorder(*122*), Schizophrenia(*123*), Sleep Duration(*124*), Tiredness(*125*), and Tobacco use disorder(*126*). We prepared summary statistics to the standard format for Linkage disequilibrium (LD) score regression. We used homologous sequences for each major cell types as a binary annotation, and the superset of all candidate regulatory peaks as the background control. For each trait, we used cell type specific (CTS) LD score regression (https://github.com/bulik/ldsc) to estimate the enrichment coefficient of each annotation jointly with the background control(*127*).

#### Fine mapping

We obtained 99% credible sets for schizophrenia from the Psychiatric Genomics Consortium website (https://www.med.unc.edu/pgc/). Potential causal variants with a posterior probabilities of association (PPA) score larger that 1% are used for overlapping with cCREs.

#### External datasets

We listed published datasets we used in this study for intersection analysis:

rDHS regions for hg38 is obtained from SCREEN database (https://screen.encodeproject.org).

The cCREs identified from adult and fetal human brains were downloaded from previously published snATAC-seq dataset.

The PhastCons^59^ conserved elements were download from the UCSC Genome Browser (http://hgdownload.cse.ucsc.edu/goldenpath/hg38/phastCons100way/).

The JASPAR motif prediction was downloaded from the UCSC Genome Browser (http://expdata.cmmt.ubc.ca/JASPAR/downloads/UCSC_tracks/2022/).

The gene expression level of Vip^+^/VIP^+^ cell type from human and mouse brain were downloaded from Allen Brain Map: Cell Types Database (https://portal.brain-map.org/atlases-and-data/rnaseq).

The gene expression level of gene *BIN1*, *MEF2A*, and *SPI1* in normal humans and patients with Alzheimer’s disease were download from Allen Brain Map: Seattle Alzheimer’s Disease Brain Cell Atlas (SEA-AD) (https://portal.brain-map.org/explore/seattle-alzheimers-disease).

#### Statistics

No statistical methods were used to predetermine sample sizes. There was no randomization of the samples, and investigators were not blinded to the specimens being investigated. However, clustering of single nuclei based on chromatin accessibility was performed in an unbiased manner, and cell types were assigned after clustering. Low-quality nuclei and potential barcode collisions were excluded from downstream analysis as outlined above.

### Supplementary Figures

**Fig. S1:**
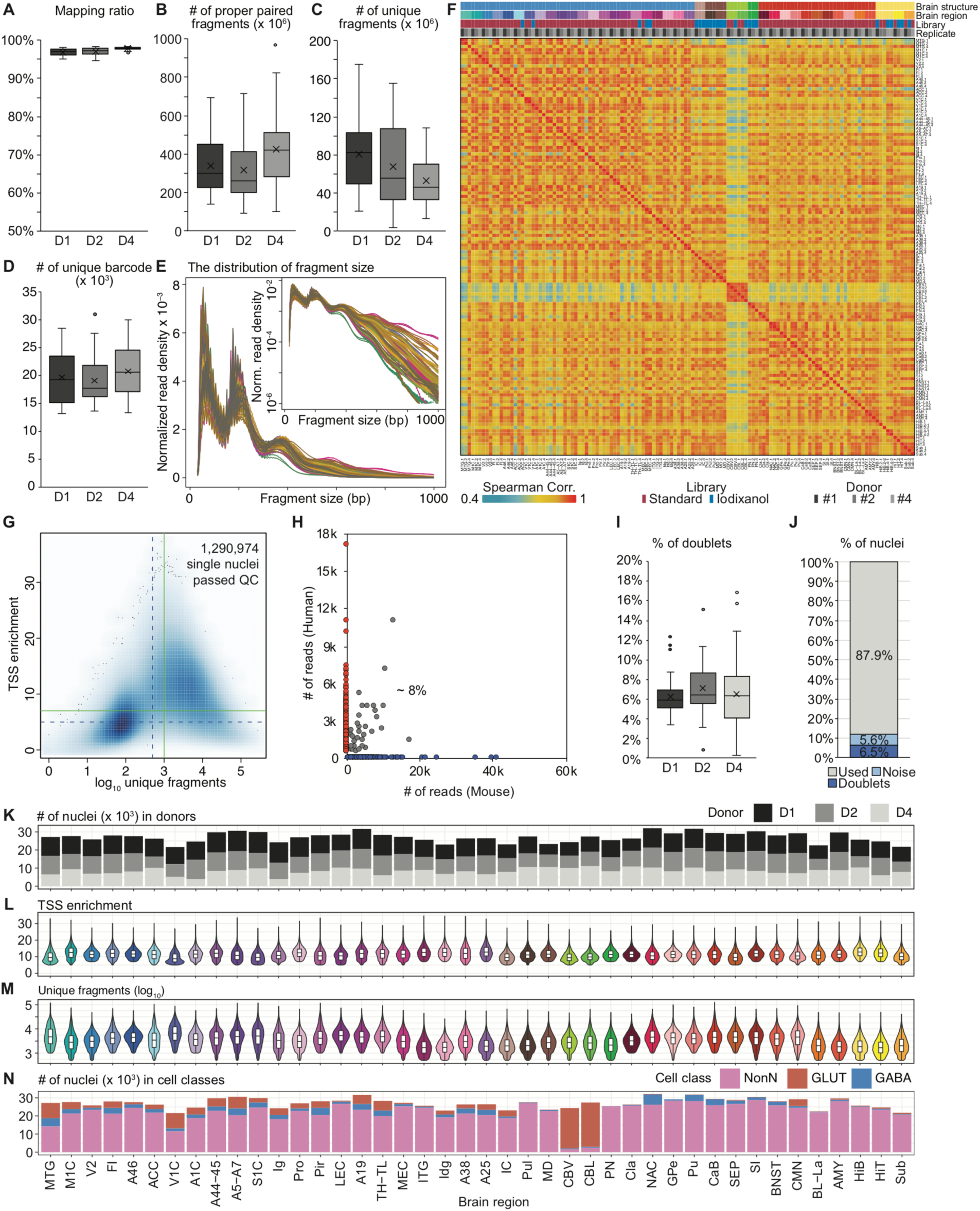
Quality control metrics of the snATAC-seq datasets. (**A**) Box plots showing the distribution of mapping ratios (the fraction of the mapped sequencing reads) in donor (biological replicate) D1, D2, and D4 of snATAC-seq experiments from each brain dissection. (**B**) Box plots showing the distribution of the number of proper read pairs (reads are correctly oriented) in donor D1, D2, and D4 of snATAC-seq experiments. (**C**) Box plots showing the distribution of numbers of unique chromatin fragments detected in donor D1, D2, and D4 of snATAC-seq experiments. (**D**) Box plots showing the distribution of number of unique barcodes captured in donor D1, D2, and D4 of snATAC-seq experiments. (**E**) Frequency distribution plot showing the fragment size distribution of each snATAC-seq dataset. (**F**) Heat map showing the pairwise Spearman correlation coefficients between snATAC-seq datasets. The column and row names consist of two parts: brain region name and donor label. (**G**) Density scatter plot illustrating fragments per nucleus and individual TSS enrichment. Nuclei in top right quadrant were selected for analysis (TSS enrichment > 5 as acceptable and >7 as ideal, and > 500 fragments per nucleus as acceptable and >1,000 fragments as ideal). (**H**) Fraction of cell collision estimated from species mix samples. (**I**) Box plot shows the fraction of potential barcode collisions detected in snATAC-seq libraries using a modified version of *Scrublet*. (**J**) Number of nuclei retained after each step of quality control. (**K**) Number of nuclei passing quality control for each brain region from donors. (**L**) Distribution of TSS enrichment. (**M**) The number of uniquely mapped fragments per nucleus for individual libraries. Boxes span the first to third quartiles (Q1 to Q3), horizontal line denotes the median, and whiskers show 1.5× the interquartile range (IQR). (**N**) Number of nuclei from three cell classes in each brain region.

**Fig. S2:**
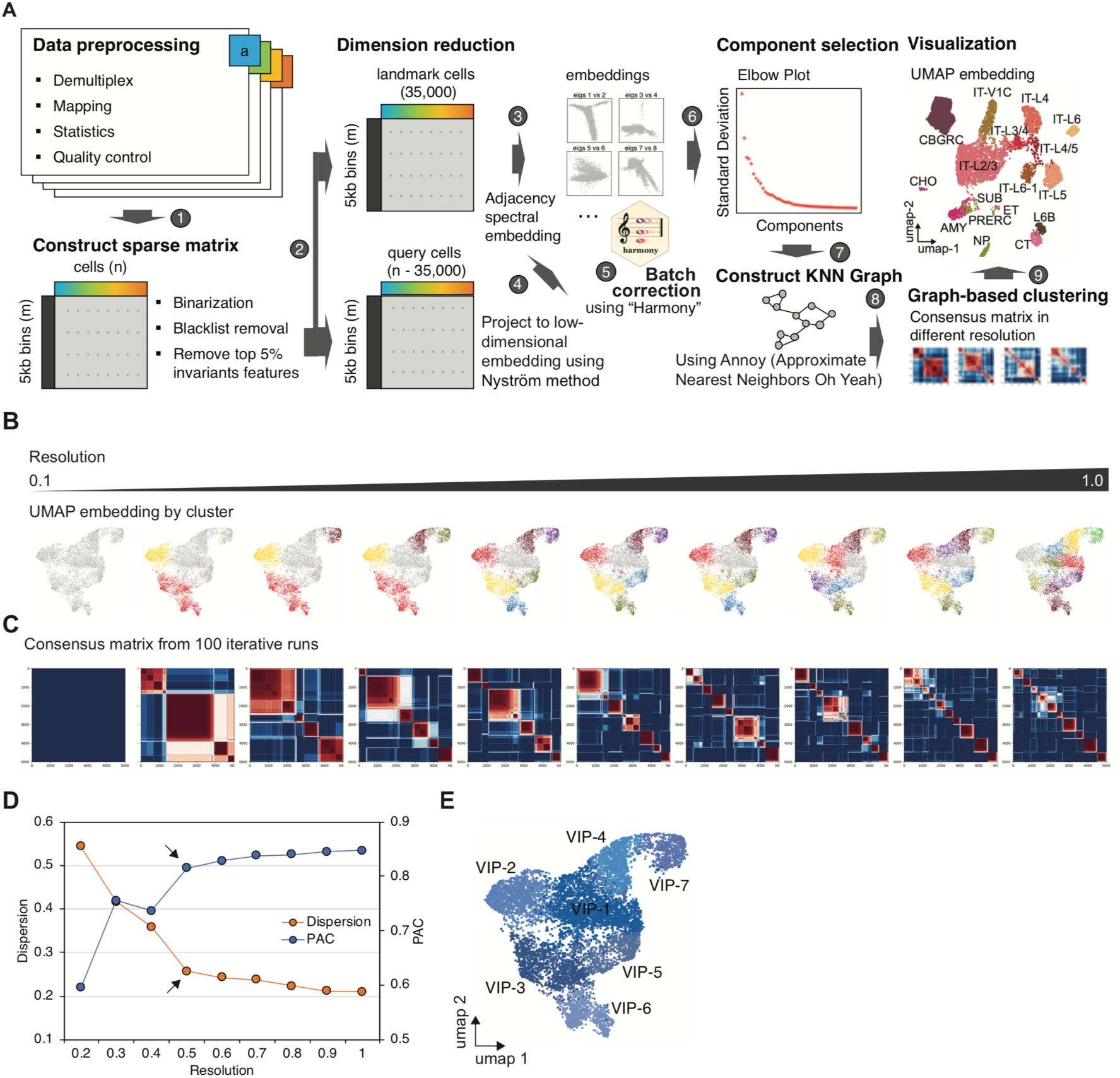
Cell clustering pipeline. (**A**) Schematic diagram of the cell clustering pipeline. (**B**) UMAP embedding of a representative cell clustering at different resolutions from 0.1 to 1.0 using Leiden algorithm. (**C**), Consensus matrix from 100 iterative clustering runs with different resolutions. (**D**) The proportion of ambiguous clustering (PAC) score and dispersion coefficient at different resolutions. A low PAC and high dispersion coefficient indicate the best and most stable clustering. (**E**) Optimal cell clustering result for a representative subclass (resolution = 0.5).

**Fig. S3:**
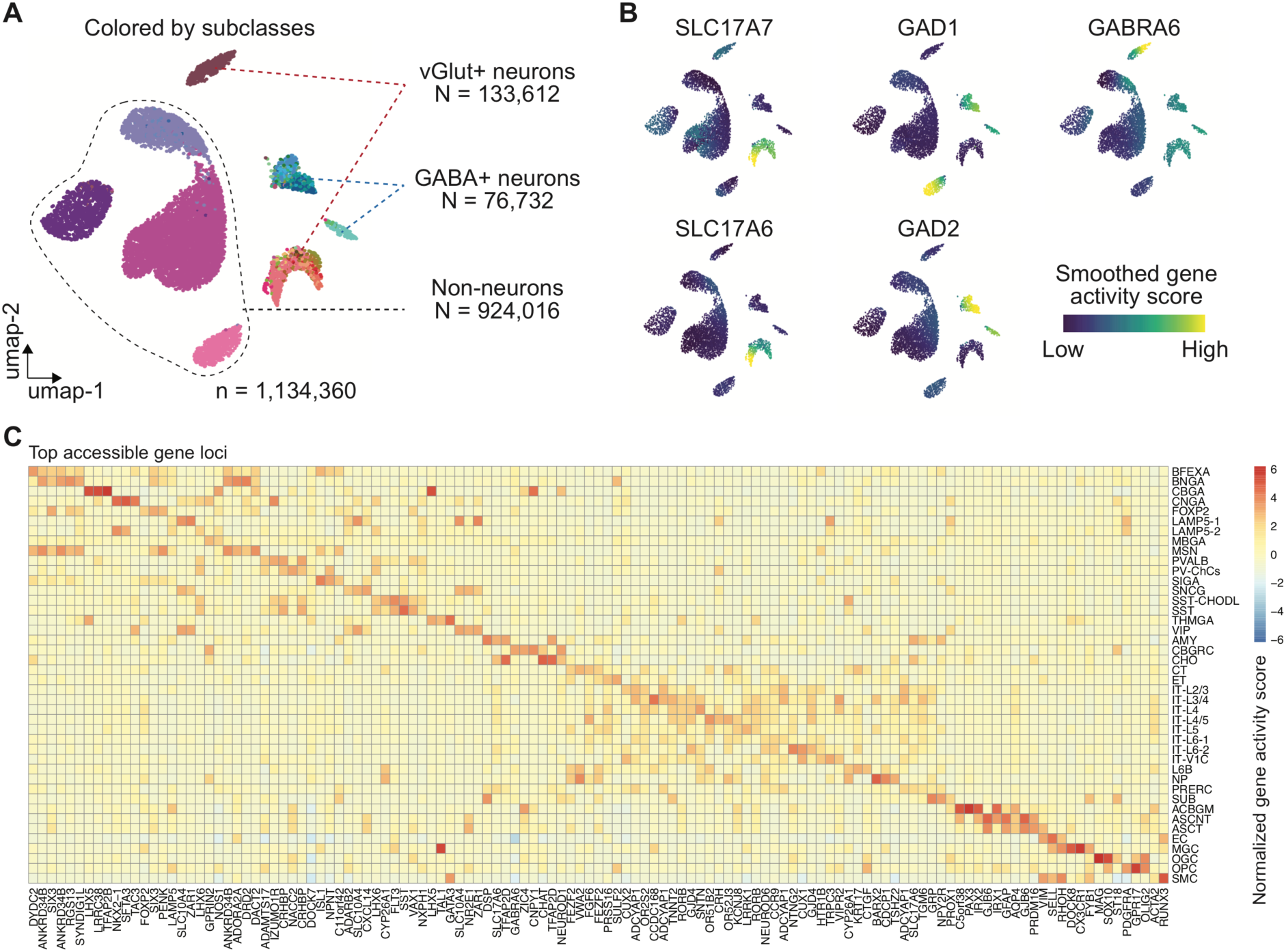
Maker genes used for annotation of different subclasses. (A) UMAP embedding of >1.1 million single nuclei and three major classes. (**B**) Smoothed gene activity score at marker loci used for annotation of three major classes. (**C**) Heat map showing normalized gene activity scores at marker gene loci used for subclass annotation.

**Fig. S4:**
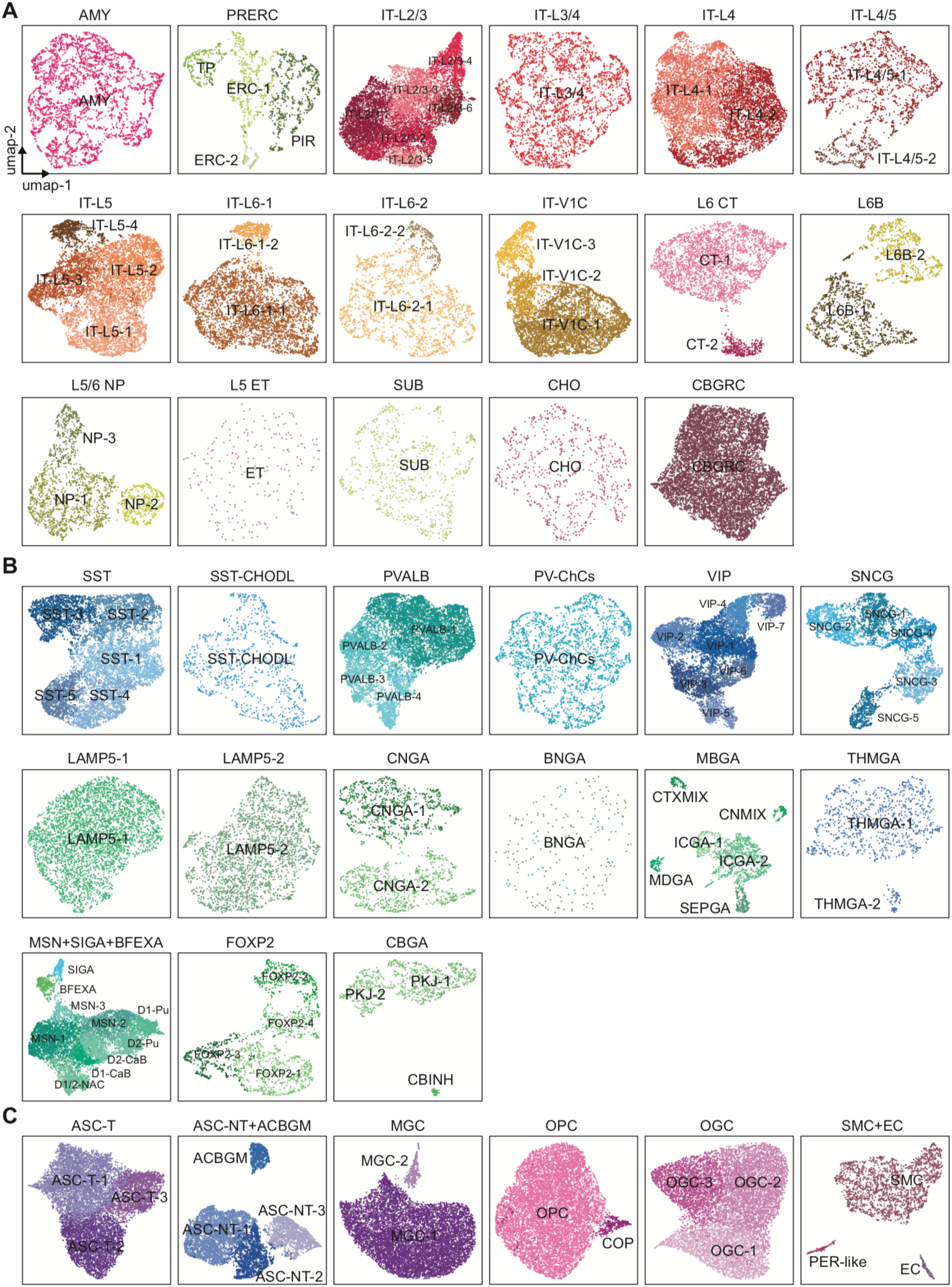
Iterative clustering identifies cell types for the subclasses. The subclasses of cerebral cells that can be further divided into cell types are shown for the glutamatergic neurons (**A**), GABAergic neurons (**B**), and non-neurons (**C**). UMAP embedding is shown for each subclass, with the subclass label shown on the top, and cell-type labels on each cell type.

**Fig. S5:**
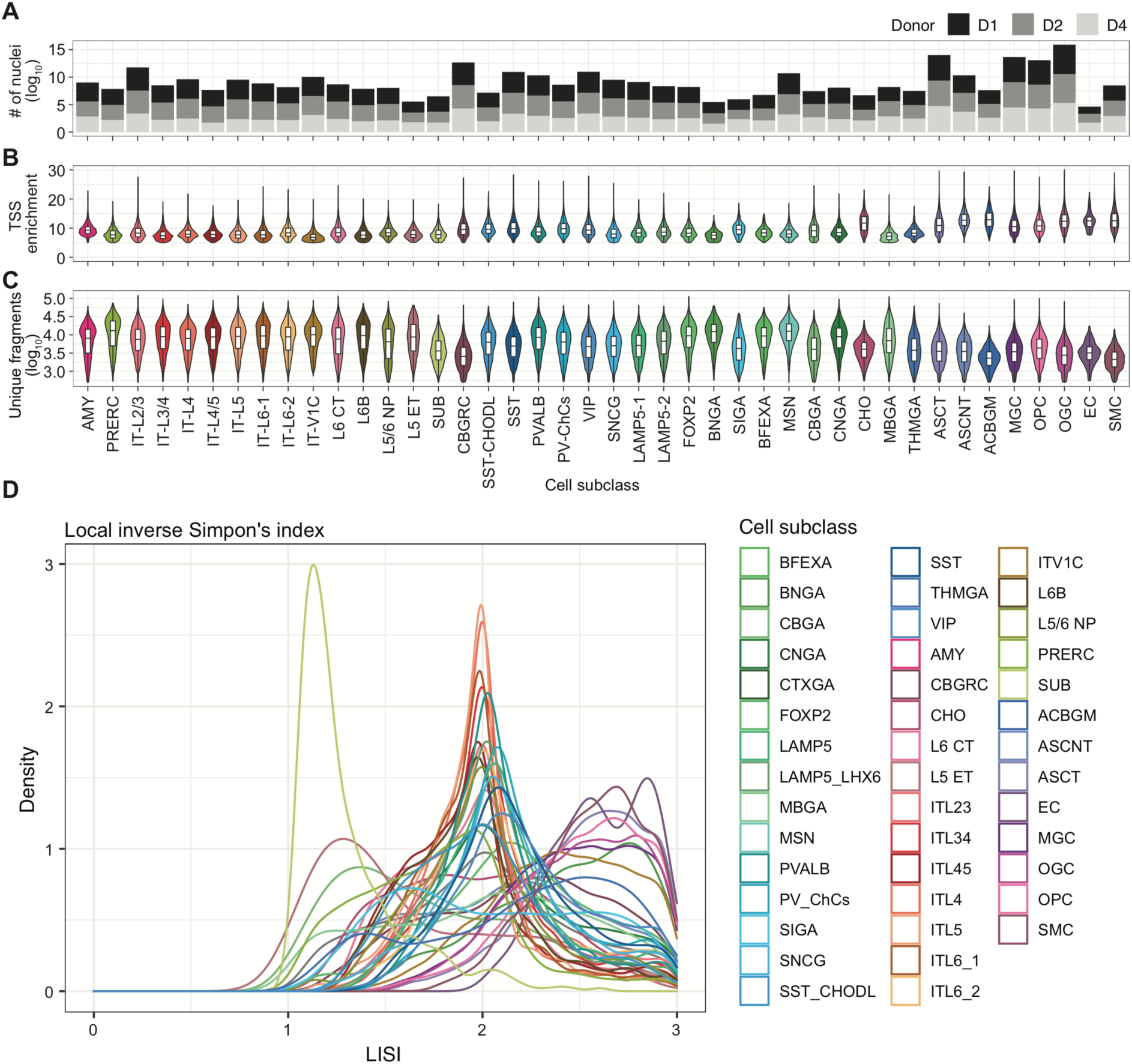
Summary statistics of cell subclasses. (**A**) Number of nuclei retained for each cell subclasses from donors. (**B**) Violin plots showing the TSS enrichment in each nucleus of each subclass. (**C**) Violin plots showing the log-transformed number of unique fragments per nucleus in each cell subclass identified. (**D**) Distribution of the local inverse Simpson’s index (LISI) scores for every single cell in each subclass.

**Fig. S6:**
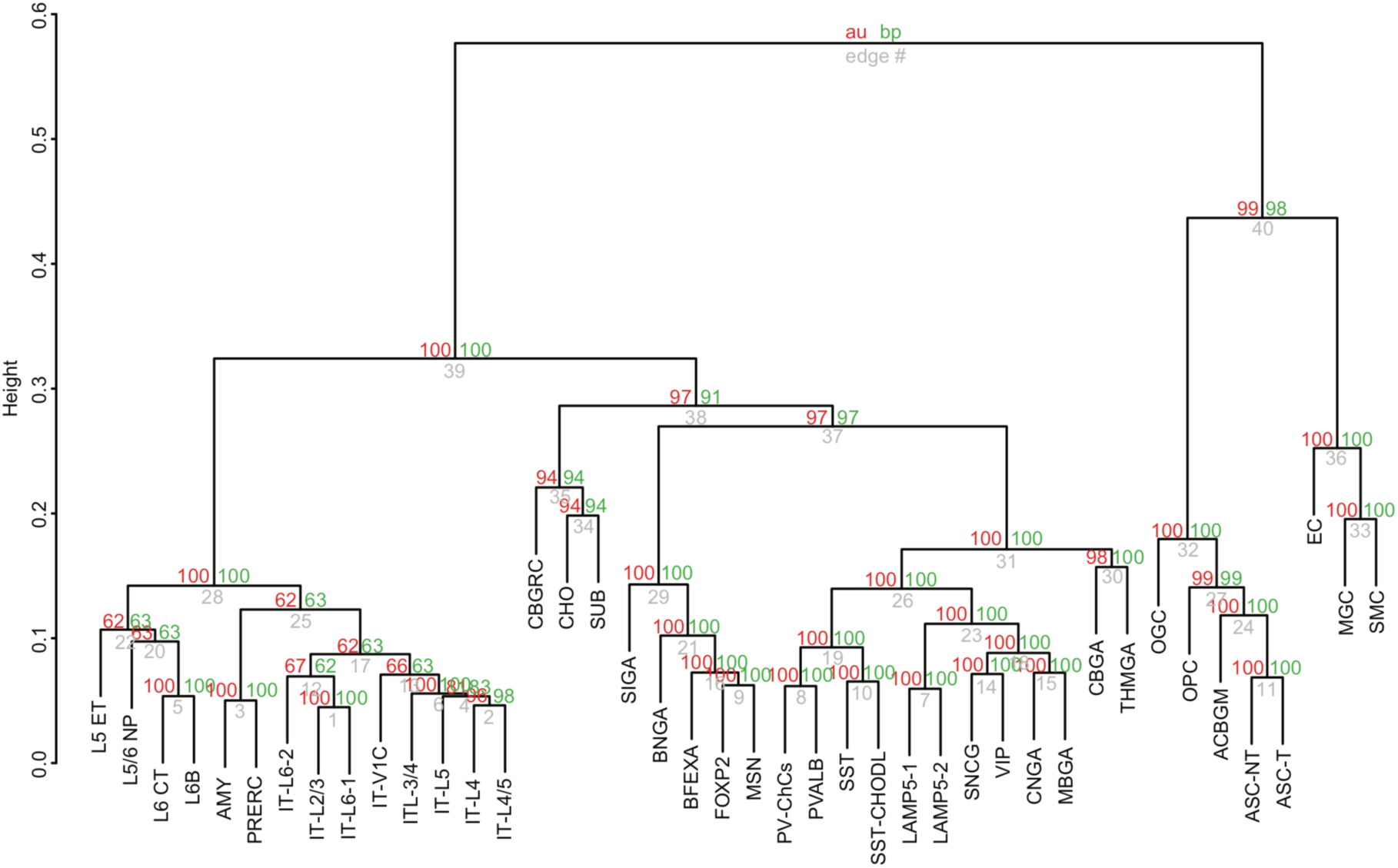
Hierarchical tree depicting the relationships between different subclasses. Dendrogram tree was constructed using 1,000 rounds of bootstrapping for the subclasses using R package *pvclust*. Nodes are labelled in grey, approximately unbiased P values (in red) and bootstrap probability values (in green) are labelled at the shoulder of the nodes, respectively. A full list and a description of cell cluster labels are in Table S4.

**Fig. S7:**
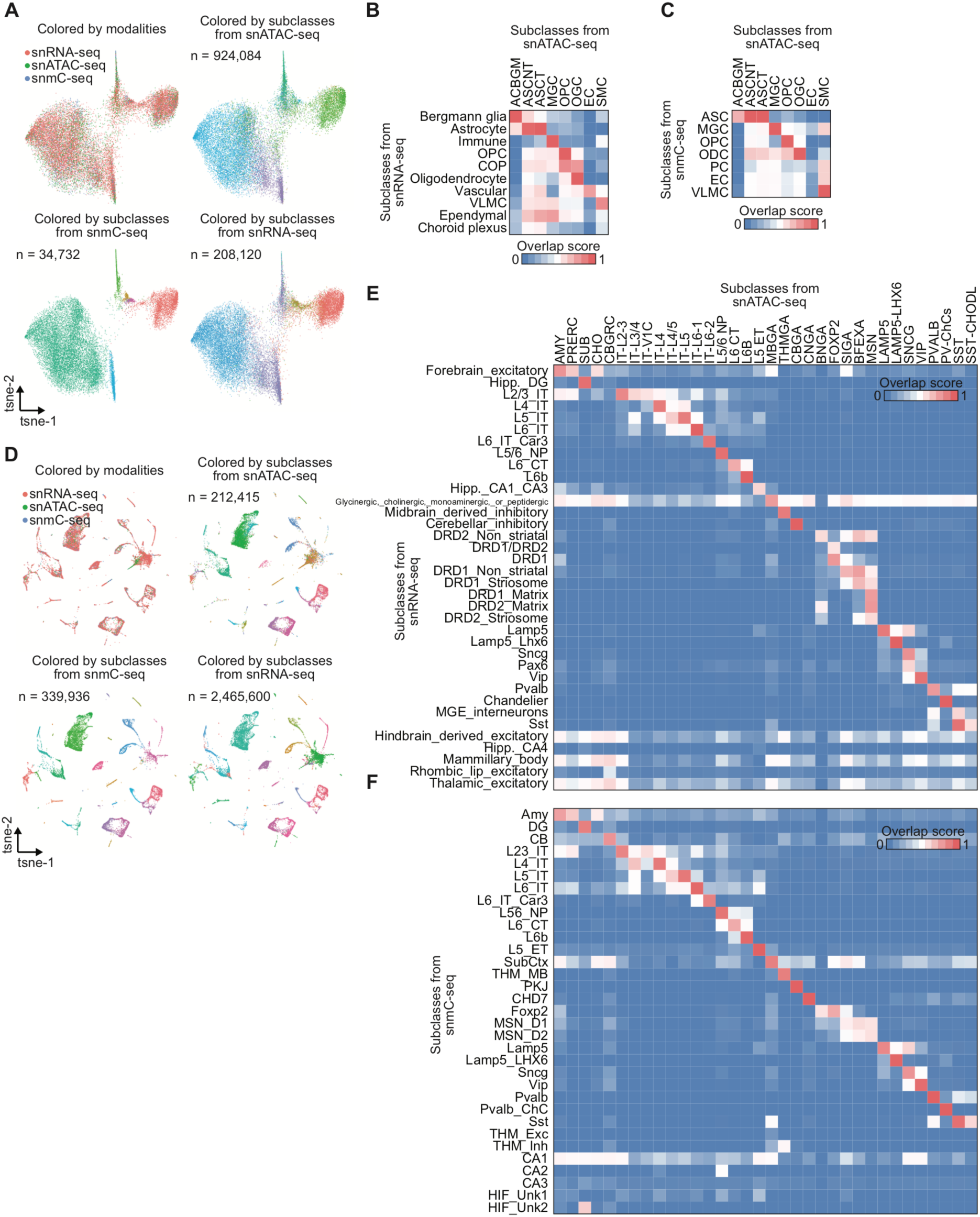
Integration analysis of snATAC-seq, snRNA-seq and snmC-seq. (**A**) UMAP embedding of non-neuronal types identified from snATAC-seq, snRNA-seq, and snmC-seq. Single cells are colored by modalities, cell subclasses defined from snATAC-seq, snmC-seq and snRNA-seq clockwise from top left. (**B**) Heat map showing the similarity between A-type (accessibility) and T-type (transcriptomics) based non-neuronal cell cluster annotations. Each row represents an A-type cluster and each column represents a T-type cluster. Similarity between original clusters and the joint cluster was calculated as the overlap score, which defined as the sum of minimal proportion of cells/nuclei in each cluster that overlapped within each co-embedding cluster. The overlap score varied from 0 to 1 and was plotted on the heatmap. For a full list of cell type labels and descriptions see Table S5. (**C**) Heat map showing the similarity between A-type (accessibility) and M-type (methylomes) based non-neuronal cell cluster annotations. (**D**) UMAP embedding of neuronal types identified from snATAC-seq, snRNA-seq, and snmC-seq. Single cells are colored by modalities, cell subclasses defined from snATAC-seq, snmC-seq and snRNA-seq clockwise from top left. (**E**) Heat map showing the similarity between A-type (accessibility) and T-type (transcriptomics) based neuronal cell cluster annotations. (**F**) Heat map showing the similarity between A-type (accessibility) and M-type (methylomes) based neuronal cell cluster annotations.

**Fig. S8:**
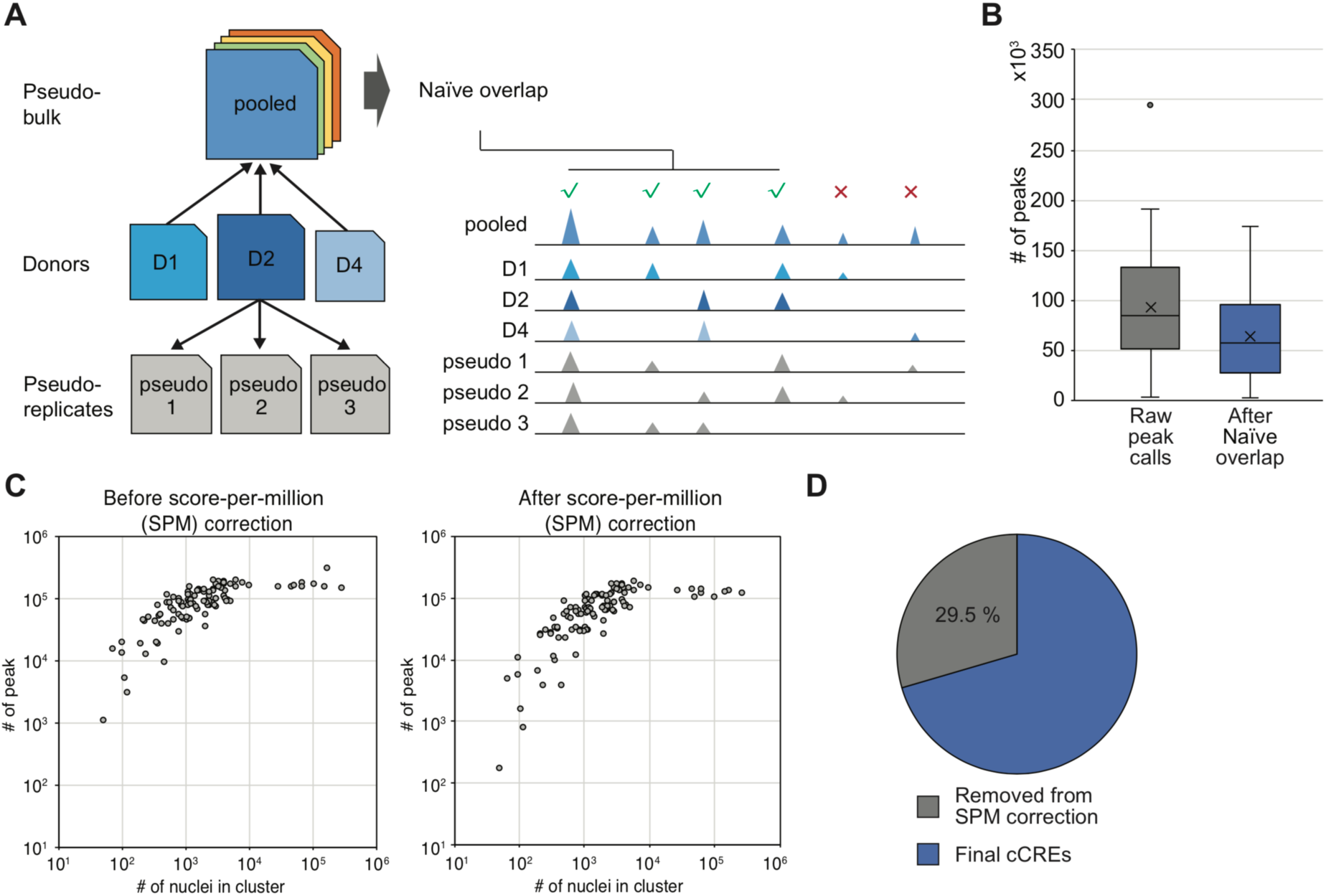
Statistics of peak calling from snATAC-seq data from each cell type. (**A**) Schematic diagram of peak calling and filtering pipeline. (**B**) Number of peaks retained after peak calling and naïve overlap filtering step for each cell type. (**C**) Scatter plots showing the relationship between the number of nuclei in each cluster and the number of peaks before score-per-million (SPM) correction (left) and after SPM correction (right). (**D**) Fraction of peaks removed in SPM correction step.

**Fig. S9:**
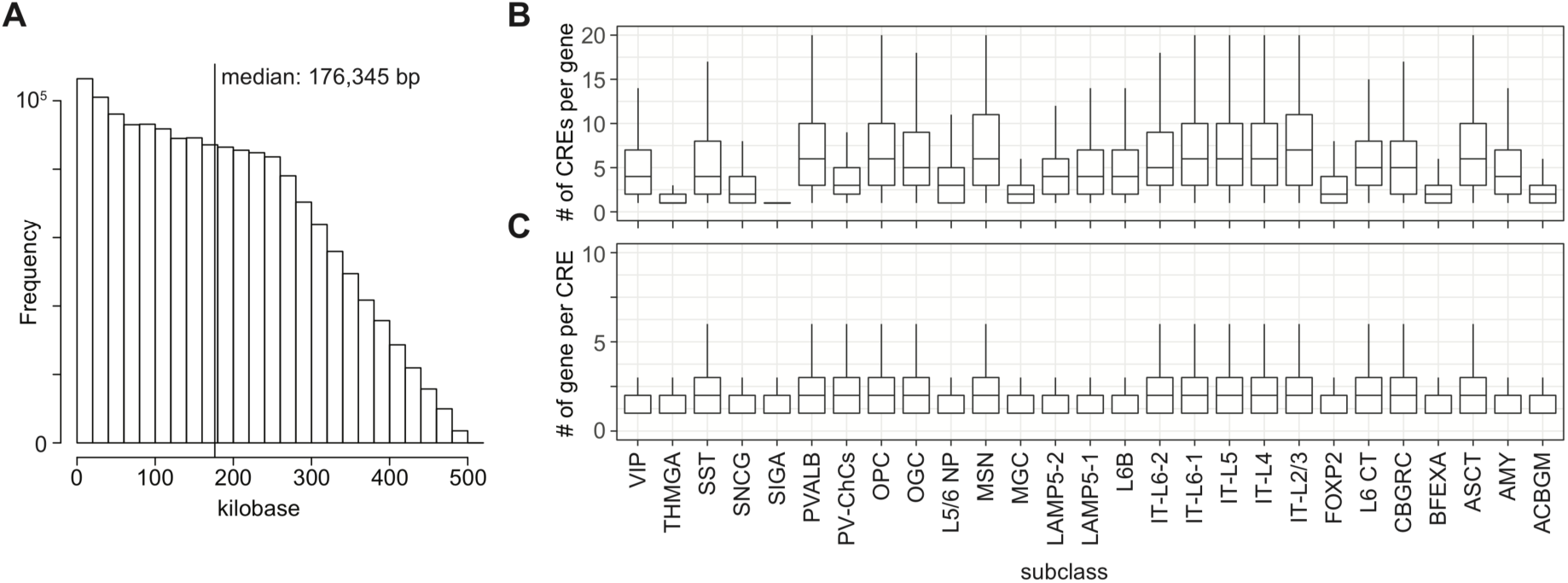
Statistics of putative enhancer-gene pairs. (**A**) Histogram illustrating the 1-D genomic distances between positively correlated distal cCRE and putative target gene promoters. (**B**) Box plot showing that genes were linked with a median of 7 putative enhancers across cell subclasses. (**C**) Box plot showing that putative were linked with a median of 2 genes across cell subclasses.

**Fig. S10:**
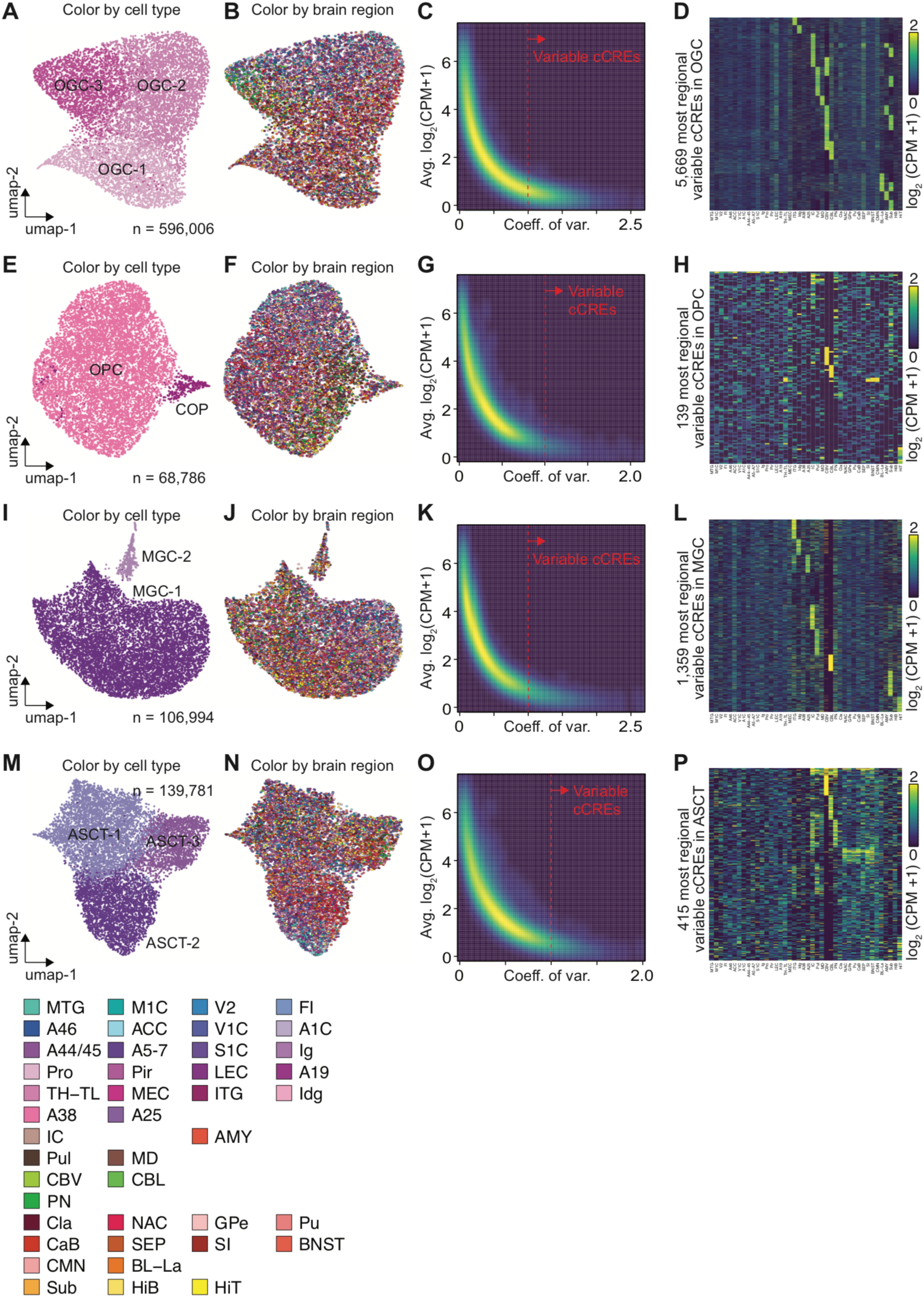
Regional specificity of glial cell types. (**A**) UMAP embedding of cell types of oligodendrocytes (OGCs). (**B**) UMAP embedding of oligodendrocytes, colored by brain regions. (**C**) Density scatter plot comparing the averaged accessibility and coefficient of variation across brain regions at each cCRE. Most variable cCREs for OGCs are defined on the right side of dash line. (**D**) Heat map showing the normalized accessibility of most variable cCREs in OGCs. CPM, counts per million. (**E**) UMAP embedding of cell types of oligodendrocyte precursor cells (OPCs). (**F**) UMAP embedding of OPCs, colored by major brain regions. (**G**) Density scatter plot comparing the averaged accessibility and coefficient of variation across brain regions at each cCRE. Most variable cCREs for OPCs are defined on the right side of dash line. (**H**) Heat map showing the normalized accessibility of most variable cCREs in OPCs. (**I**) UMAP embedding of cell types of microglia (MGC). (**J**) UMAP embedding of MGC, colored by major brain regions. (**K**) Density scatter plot comparing the averaged accessibility and coefficient of variation across brain regions at each cCRE. Most variable cCREs for MGC are defined on the right side of dash line. (**L**) Heat map showing the normalized accessibility of most variable cCREs in MGC. (**M**) UMAP embedding of cell types of astrocytes (ASCs). (**N**) UMAP embedding of ASCs, colored by major brain regions. (**O**) Density scatter plot comparing the averaged accessibility and coefficient of variation across brain regions at each cCRE. Most variable cCREs for ASCs are defined on the right side of dash line. (**P**) Heat map showing the normalized accessibility of most variable cCREs in ASCs.

**Fig. S11:**
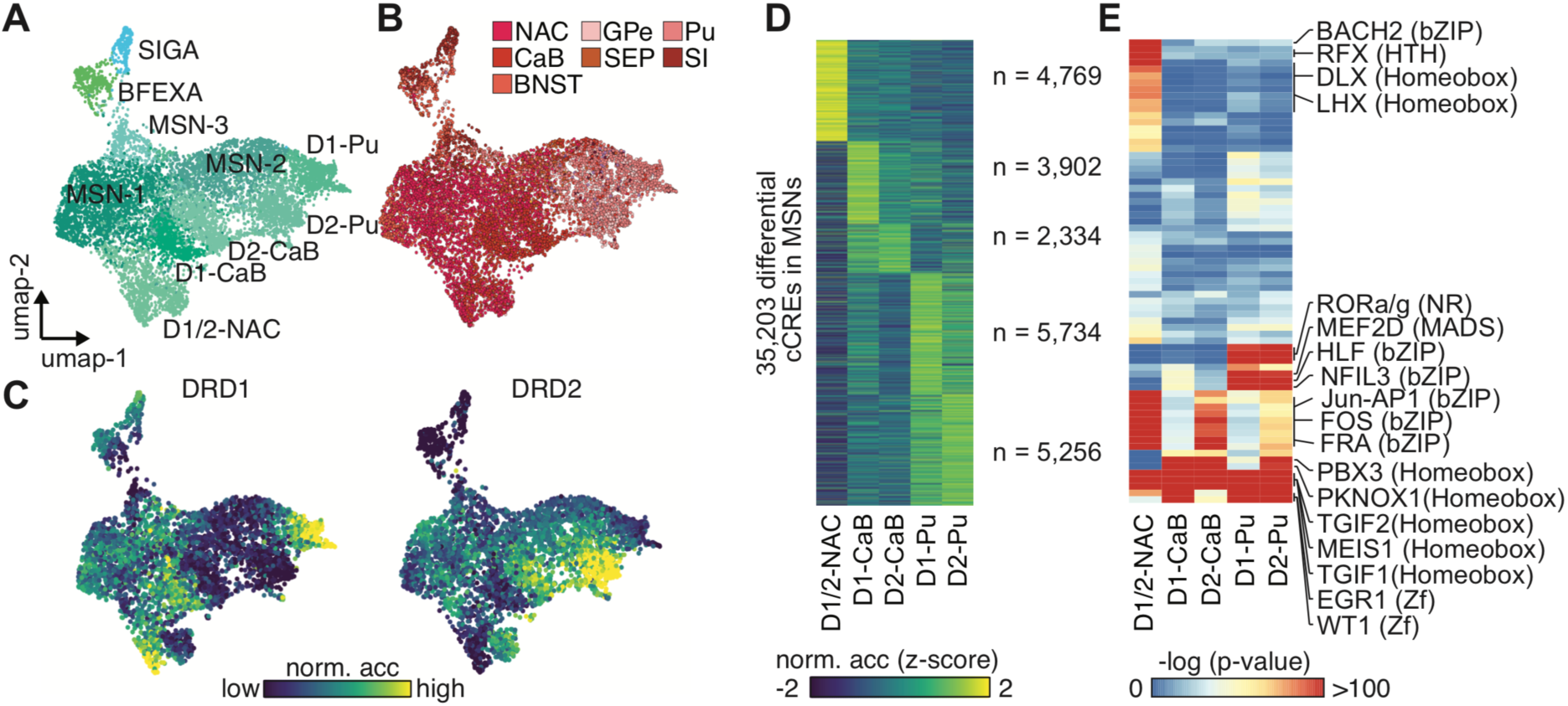
Regional difference in MSN. (**A**) UMAP embedding of cell types of medium spiny neurons (MSNs) and cell types from cerebral nuclei. (**B**) UMAP embedding of MSNs, colored by brain regions. (**C**) UMAP embedding of normalized accessibility at gene *DRD1* and *DRD2* in MSNs. (**D**) Normalized chromatin accessibility of 35,203 MSN cell-type-specific cCREs. (**E**) Enrichment of known transcription factor (TF) motifs in MSN cell-type-specific cCREs.

**Fig. S12:**
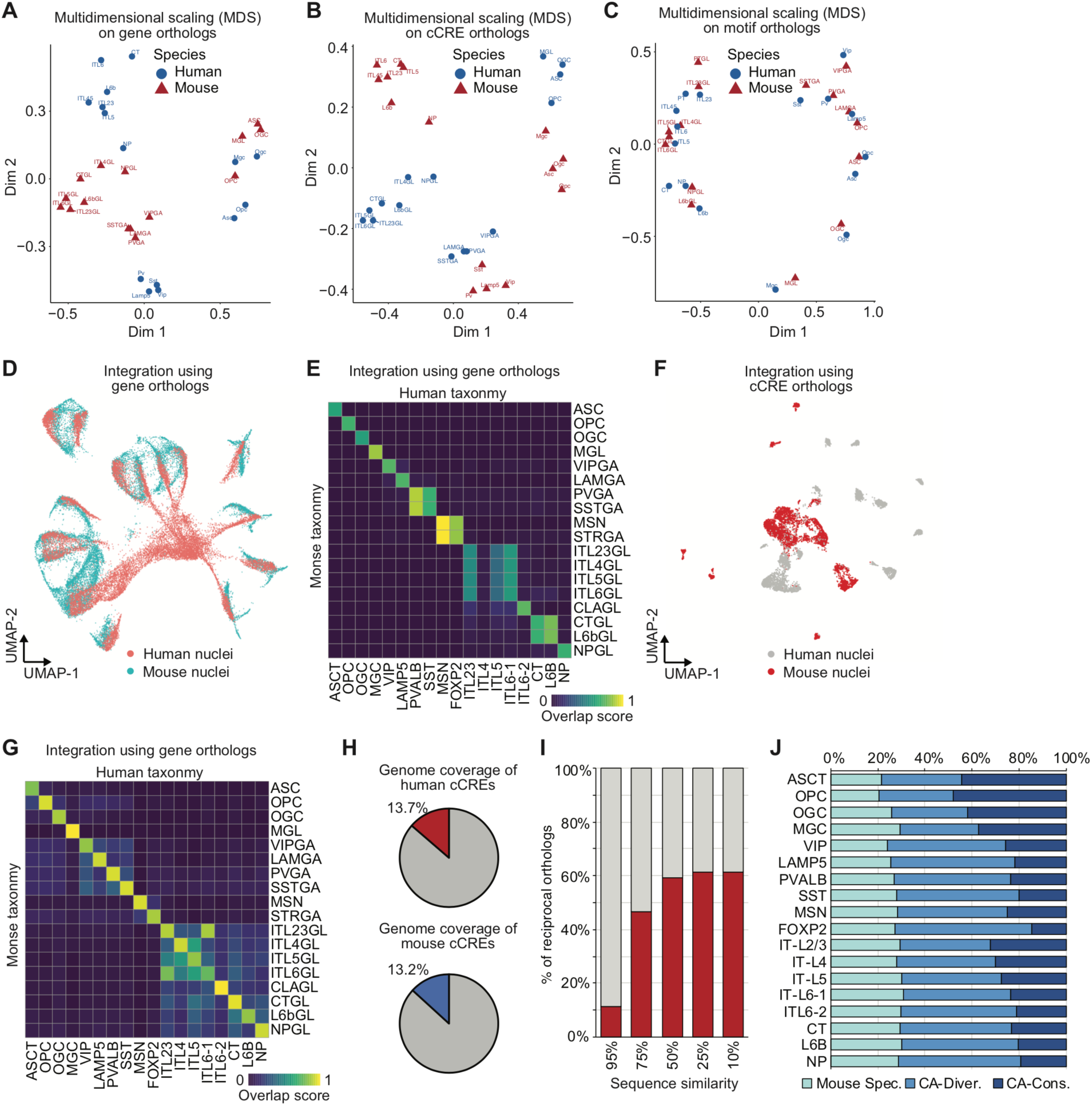
Integration analysis of snATAC-seq data between human and mouse cerebrum. (**A**) Multidimensional scaling (MDS) embedding of human and mouse subclasses using gene orthologs. (**B**) MDS embedding of human and mouse subclasses using cCRE orthologs. (**C**) MDS embedding of human and mouse subclasses using motif orthologs. (**D**) Co-embedding analysis of single nuclei from human and mouse subclasses using “gene activity score” from gene orthologs. (**E**) Heat map illustrating the overlap between cell subclasses from both species from integration analysis using gene orthologs. Rows show cell subclasses from mouse cerebrum; columns show cell subclasses from the human cerebrum. The overlap between the human and mouse cell subclasses was calculated (overlap score) and plotted on the heat map. (**F**) Co-embedding analysis of single nuclei from human and mouse subclasses using chromatin accessibility from cCRE orthologs. (**G**) Heat map illustrating the overlap between cell subclasses from both species from integration analysis using motif orthologs. (**H**) Genome coverage of cCREs from used human and mouse subclasses. (**I**) Percentage of reciprocal orthologous cCREs identified in human by using different cutoff of sequence similarity. (**J**) bar plot showing three categories of mouse cCREs in corresponding cell subclasses.

**Fig. S13:**
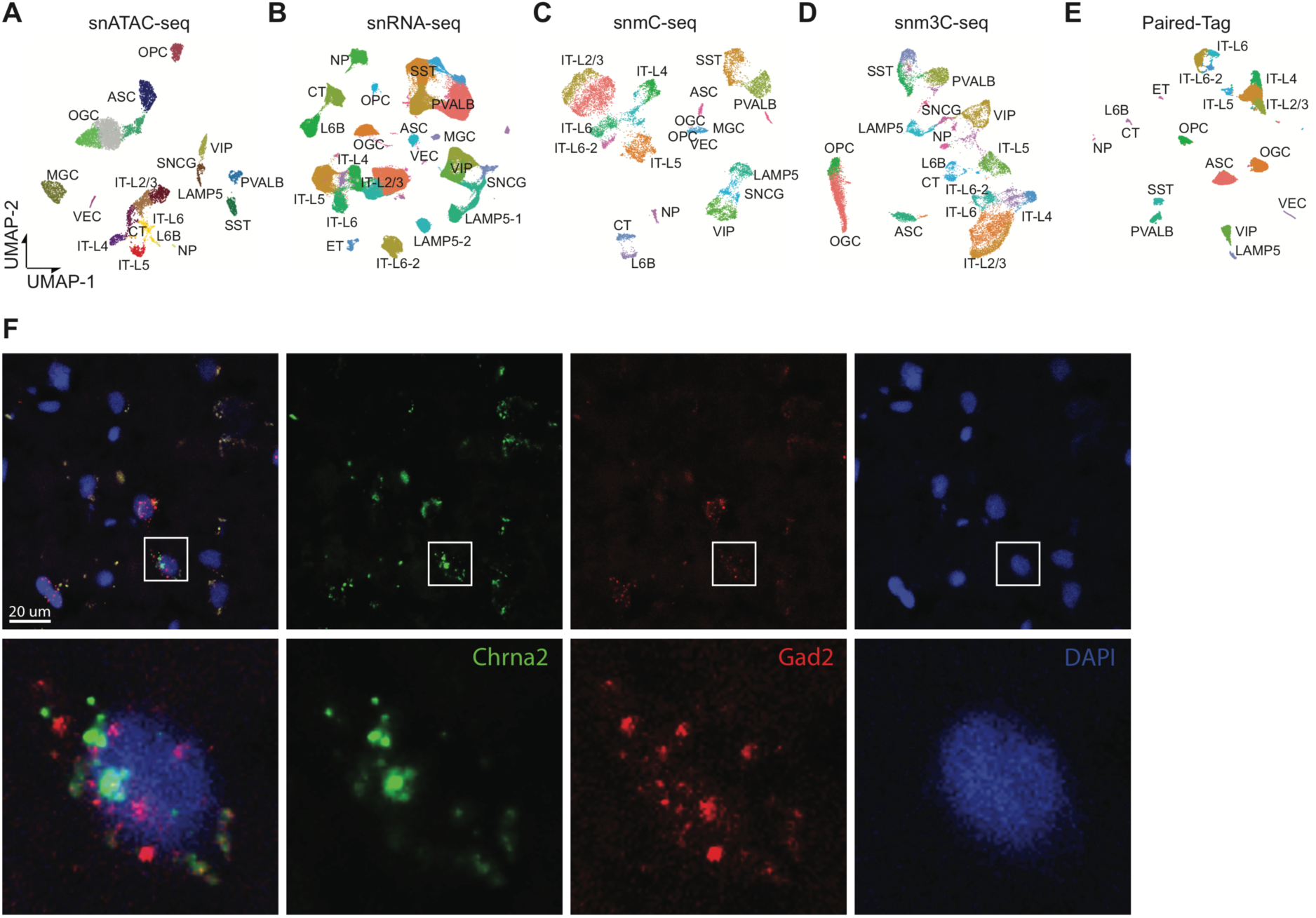
Integration analysis of multi-modal omics datasets. UMAP embedding of cell clustering and annotation for snATAC-seq (**A**), snRNA-seq (**B**), snmC-seq (**C**), snm3C-seq (**D**), Paired-Tag (**E**) from brain region M1C and MTG. (**F**) RNAscope validation of *CHRNA2* position lineage of VIP^+^ GABAergic neurons.

**Fig. S14:**
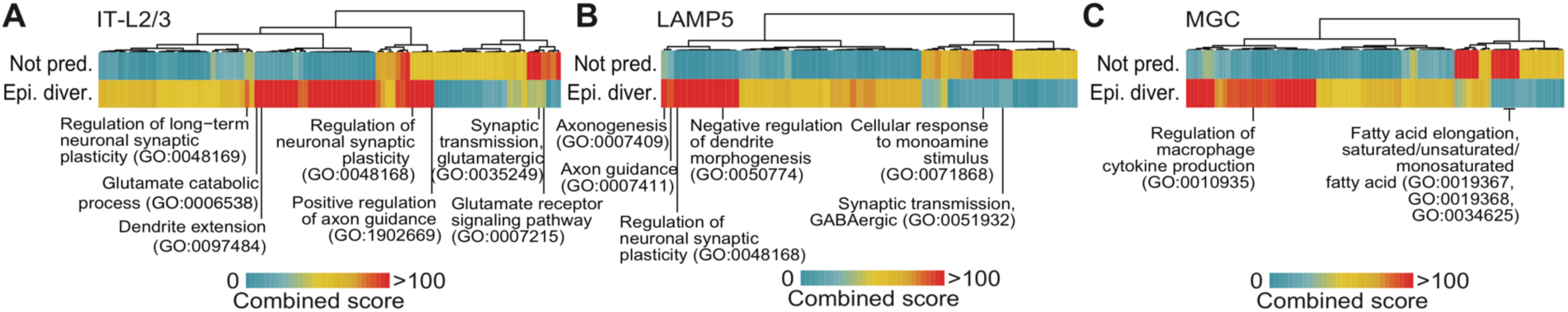
Gene ontology of putative target genes that linked with human divergent cCREs predicted and not predicted from mouse gkmsvm models. (**A**) Gene ontology of putative target genes that linked to human divergent cCREs in IT-L2/3, which can be predicted and cannot predicted from mouse gkmsvm model. (**B**) Gene ontology of putative target genes that linked to human divergent cCREs in IT-L2/3, which can be predicted and cannot predicted from mouse gkmsvm model. (**C**) Gene ontology of putative target genes that linked to human divergent cCREs in IT-L2/3, which can be predicted and cannot predicted from mouse gkmsvm model. The combined score is defined in R package *Enrichr*, which calculated by multiplying the log-transformed p-value from Fisher’s exact test with the z-score computed to assess the deviation from the expected rank.

**Fig. S15:**
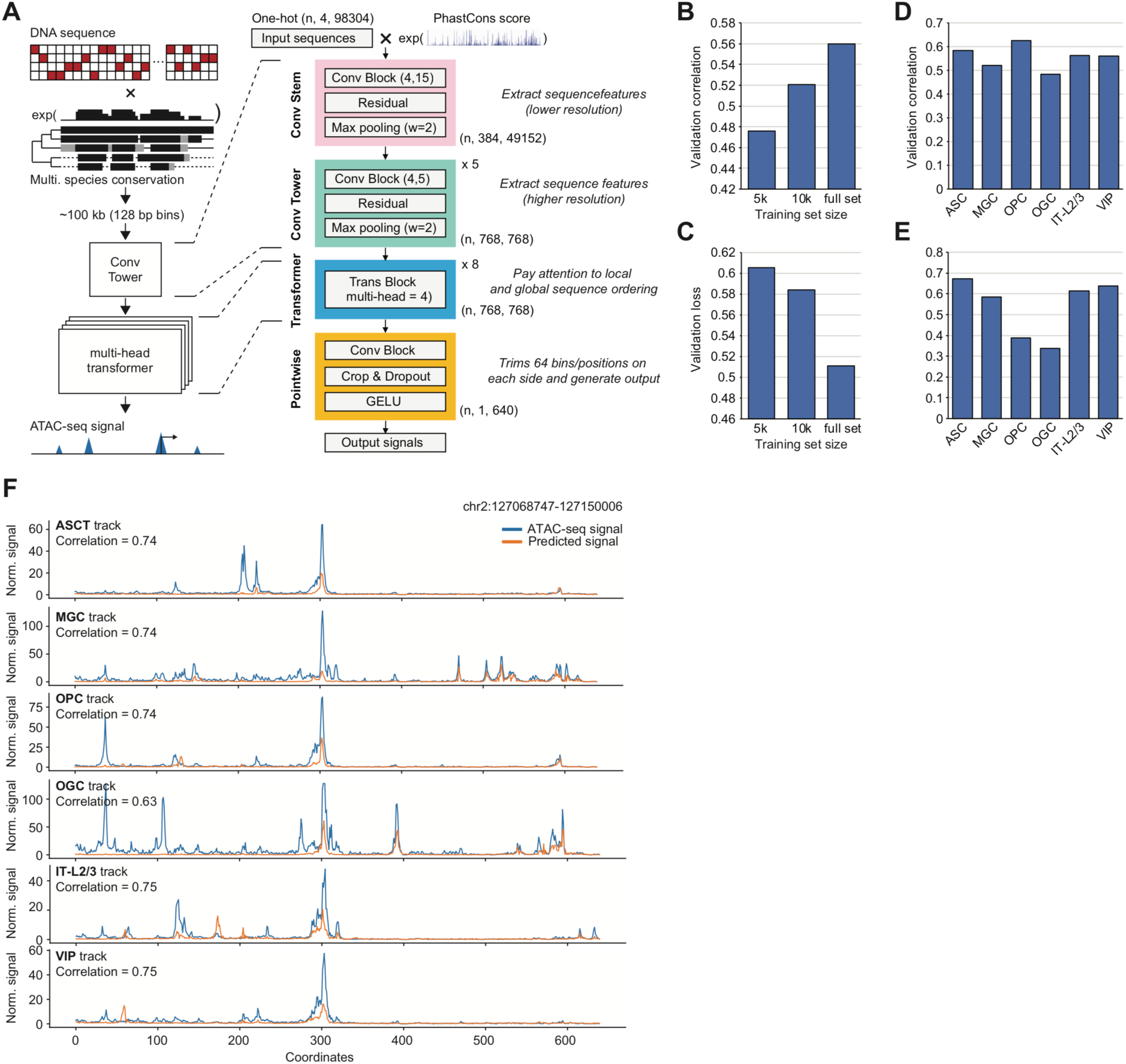
Deep learning model architecture and statistics. (**A**) The graphic diagram showing the model architecture. Output shapes (without batch dimensions) are shown as tuples on the right side of the blocks. n, number of inputs in mini-batch, w, width for padding. (**B**) Benchmark of model performance on validation set using different size of training set. The Pearson correlation coefficient showed was calculated between true ATAC-seq signals and predicted signals. (**C**) Benchmark of loss on validation set using different size of training set. The loss showed was calculated from function *PoissonNLLLoss*. (**D**) Bar plots showing model performance on different human cell subclasses. (**E**) Bar plots showing model loss on different human cell subclasses. (**F**) Predicted signal centered at BIN1 promoter from models trained from different cell subclasses.

### Supplementary Tables

**Table S1:** Summary of brain samples and dissections.

**Table S2:** Metadata for the snATAC-seq experiments.

**Table S3:** Metatable and annotation of single nuclei.

**Table S4:** Human brain cell taxonmy and annotation.

**Table S5:** Cell subclass annotation and marker genes.

**Table S6:** List of cCREs defined in the current study.

**Table S7:** Cell type assignment of cCREs.

**Table S8:** Genomic annotation of cCREs defined in the current study.

**Table S9:** The cis regulatory modules and the cell subclasses

**Table S10:** The cis regulatory modules and cCREs.

**Table S11:** Known motif enrichment in the cis regulatory modules.

**Table S12:** Summary of gene-cCRE correlations.

**Table S13:** Association modules of enhancer-gene pairs with RNA-ATAC matched cell subclasses.

**Table S14:** Association modules of enhancer-gene pairs with individual putative enhancers.

**Table S15:** Known motif enrichment in the putative enhancers from different modules.

**Table S16:** Differential cCRE in the cell types of astrocyte.

**Table S17:** Known motif enrichment in the cell types of astrocyte.

**Table S18:** Differential cCRE in the cell types of D1-/D2-MSN.

**Table S19:** Known motif enrichment in the cell types of D1-/D2-MSN.

**Table S20:** List of different categories of cCREs.

**Table S21:** Genomic annotation of different categories of cCREs.

**Table S22:** Differential cCRE in the cell types of VIP^+^ neurons.

**Table S23:** Different categories of cCREs marked by K27ac and K27me3.

**Table S24:** Gene Ontology analysis of the putative target gene of different categories of cCREs.

**Table S25:** List and references of genome-wide association studies.

**Table S26:** LDSC analysis of cCREs for every cell type.

**Table S27:** LDSC analysis of different categories of cCREs.

**Table S28:** in silico mutagenesis on microglia-specific enhancer.

**Table S29:** Primer sequences and nuclei barcodes.

## Notes

http://catlas.org

http://catlas.org/humanbrain

